# Dual-loop involving microbial single-cell protein production from soybean-processing wastewater and effluent-based refinement for circular bioeconomy applications

**DOI:** 10.64898/2026.07.08.737151

**Authors:** Ramanujam Srinivasan Vethathirri, Ezequiel Santillan, Chia Chee Ng, Stefan Wuertz

**Affiliations:** Singapore Centre for Environmental Life Sciences Engineering, Nanyang Technological University, Singapore, 637551, Singapore; School of Civil and Environmental Engineering, Nanyang Technological University, Singapore, 639798, Singapore

**Keywords:** soybean wash water, inoculated microbial consortia, hydraulic retention time, microbial cell protein, dual-loop bioprocess, downstream valorisation, circular bioeconomy

## Abstract

Nutrient-rich food-processing wastewaters represent valuable yet under-utilised side streams for sustainable protein production in the form of microbial biomass. Here we present an integrated dual-loop bioprocess that converts soybean-processing wastewater into microbial single-cell protein (SCP) while achieving substantial nutrient removal and product refinement. In the first loop, previously enriched microbial consortia were inoculated and cultivated in four parallel sequencing batch reactors (SBRs) for 44 days at a hydraulic retention time (HRT) of 3 days. This bioprocess configuration demonstrated features that support future scale-up while maintaining process stability, achieving a protein content of 33.3L±L3.2%, doubling the protein yield (15.32L±L3.49Lg dry weight per g soluble TKN) and quadrupling the production rate (0.29L±L0.06Lg dry weight LL¹ dL¹) compared to operating reactors without inoculation (HRT: 7.2 days). Effluent treatment was stable, with 84% carbon and 78% nitrogen removal efficiencies, demonstrating efficient nutrient recovery. The SCP biomass was enriched in functional taxa, including *Acidipropionibacterium*, *Lactococcus*, *Megasphaera,* and *Azospirillum*, suggesting that reactor conditions and inoculum selection promoted a stable, protein-productive microbial community with potential probiotic benefits. In the second loop, bioreactor effluent was reused as aqueous matrix for heat treatment (60°C) of the SCP biomass, reducing the RNA content from 8.6% to 2.6%, with a 39% biomass loss accompanied by a 30% increase in total amino acid concentration. Hence, our valorisation approach integrates microbial biomass production, effluent reuse, and product refinement within a circular framework. The system provides a resource-efficient pathway for converting food-sector side streams into high-quality microbial community-based SCP, highlighting its potential scalability for sustainable nutrient and water management.

## 1. Introduction

The urgent global challenges of ensuring food security, sustainable water management, and environmental stewardship are central to the United Nations Sustainable Development Goals (SDGs) (Pérez-Escamilla, 2017). The ever increasing protein demand driven by population growth and changing diets exerts substantial pressure on conventional protein sources, such as livestock and fisheries (Henchion et al., 2017; Hu et al., 2024), and contributes significantly to land degradation, water scarcity, and greenhouse gas emissions (Adetola et al., 2025).

Microbial community-based protein produced from diverse microbial consortia cultivated on nutrient-rich effluents produced in the food and beverage sector offers a sustainable, scalable solution (Vethathirri et al., 2021; Thi et al., 2025) that also benefits the implementation of a circular bioeconomy (Durkin et al., 2022). This approach not only addresses protein deficiency with minimal resource inputs but also valorises industrial wastewaters, reducing waste and environmental pollution. Furthermore, single-cell protein (SCP) production supports climate mitigation by lowering reliance on traditional agriculture (Matassa et al., 2016), alleviates pressure on marine ecosystems through sustainable aquafeed alternatives (Hua et al., 2019; Pilmer et al., 2025), and advances innovative bioprocessing technologies. Collectively, these integrated benefits position microbial protein production as a pivotal strategy to contribute to multiple SDGs simultaneously.

Yield (g SCP/g substrate) and production rate (g SCP/L/d) are two metrics commonly used for evaluating microbial community-based protein production, providing insights into the efficiency and scalability of the process (Cao et al., 2021; Peng et al., 2022; Sørensen et al., 2025). The performance of SCP production is influenced by various operating parameters, including pH, solids retention time (SRT), hydraulic retention time (HRT), cycle time, aerobic-to-anoxic phase ratio, mixing rate, and temperature (Vethathirri et al., 2021). Batch systems, particularly sequencing batch reactors, offer an effective platform for testing these conditions, as they integrate anaerobic and aerobic phases within a single cycle, allowing for flexible adjustments to optimize production (Clagnan et al., 2024). Despite the advantages of these systems, there remains a lack of comprehensive data to define an optimal range of operating conditions for SCP production, particularly when considering the specific characteristics of influent wastewater, which can vary significantly and impact microbial community performance.

Controlling the retention time of the substrate by adjusting the HRT is a key operational strategy for optimizing single-cell protein (SCP) production. Previous studies have shown that the HRT can influence intracellular storage components such as polyhydroxyalkanoates (Shang et al., 2025) and affect protein accumulation in microbial biomass, although the reported effects vary across systems and feedstocks (Muys et al., 2020). These contrasting outcomes highlight the need to better understand how the HRT influences SCP yield, production rate, and effluent quality, particularly for variable food-processing wastewaters. In an earlier study, reactors operated at an HRT of 7.2 days without inoculation achieved stable operation but moderate yields (Vethathirri et al., 2025). Hence, a shorter HRT combined with inoculation with a previously enriched microbial community may offer a strategy to improve productivity while maintaining wastewater treatment performance. By evaluating these parameters together with biomass yield, production rate, and nutrient removal efficiency, we aim to design efficient and circular SCP bioprocesses based on nutrient-rich wastewaters produced in the food and beverage industry.

Ribonucleic acid (RNA) content in microbial biomass is a key nutritional and regulatory consideration in developing single-cell protein (SCP) for feed applications. Due to rapid growth and high metabolic activity, microbial cells accumulate substantial nucleic acids alongside protein, particularly in bacterial SCP where nitrogen is incorporated into amino acids and nucleotides (Sharif et al., 2021). In humans and other primates lacking urate oxidase, excessive intake of RNA-rich food can elevate uric acid and lead to gout or kidney disorders (Cheng et al., 2023; Roman, 2023). Consequently, SCP for human or pet consumption often undergoes RNA reduction (Ritala et al., 2017). In aquaculture, however, the relevance of RNA as a limiting factor remains debated. Fish species such as sea bass and salmonids possess functional uricase systems that metabolize purine-derived uric acid efficiently (Li & Gatlin., 2006), and low dietary nucleic acid levels (0.5–0.75 g kgL¹ feed) can enhance immunity (Rairat et al., 2022). These findings suggest that RNA is not inherently anti-nutritional. Given the current limited evidence and apparent species-specific effects of RNA-containing diets on aquaculture animals, there is a need to understand the consequences of removing RNA in alternative protein formulations. We hypothesized that lowering the RNA content in SCP would improve amino acid bioavailability.

The objective of this study is to assess how the combined application of inoculation and a shortened HRT influences microbial community–based SCP production from soybean-processing wastewater. In addition, we aimed to determine whether the resulting SCP can be further refined by reducing its RNA content using the effluent from the SCP-producing bioreactor. These research questions guided the development of our dual-loop framework, in which food-grade side stream is first converted into microbial biomass and the subsequent effluent is repurposed for RNA reduction when such refinement is desired (Figure 1).

**Figure 1.**
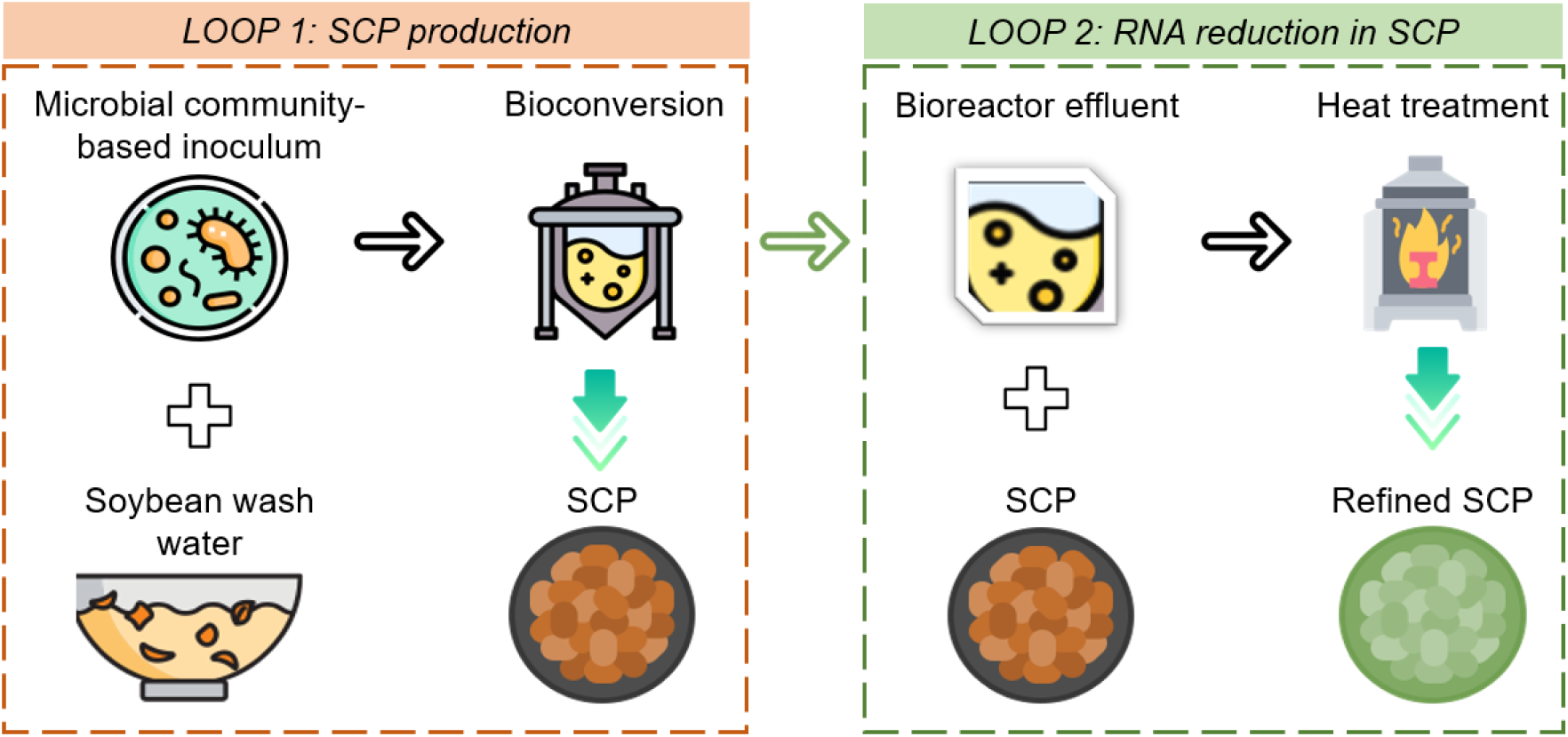
Conceptual dual-loop schematic illustrating the integrated process for microbial protein production from soybean-processing wastewater. Loop 1 targets high-yield, high-productivity SCP generation using a defined microbial inoculum and short hydraulic retention time (3 days), while maintaining effluent quality. Loop 2 repurposes the process effluent for a downstream refinement stage aimed at RNA reduction and biomass compositional enhancement. This dual-loop design exemplifies circular resource use by integrating both valorisation and internal upgrading within a closed bioprocessing system.

## 2. Materials and methods

### 2.1 Experimental design

Four 4-L bioreactors were operated as sequencing batch reactors (SBRs) for 44 days under identical conditions with intermittent aeration, using wastewater obtained from a soybean processing company in Singapore. Over two months, fourteen 20-L carboys of soybean-soaking wastewater were collected on six occasions (see e-Supplementary Materials, Table S1). Samples were taken from each reactor three times per week on alternate days to assess weekly variations in the measured parameters.

### 2.2 Operational parameters and bioreactor arrangement

The reactor working volume was 4 L, with an additional headspace of approximately 2 L. The reactor temperature was maintained at 30 °C and sludge was continuously mixed at 375 rpm. The feeding phase occurred during the initial 10 min of a cycle, followed by 180-min anoxic/anaerobic and 1190-min aerobic phases. Once the cycle had finished according to a feeding scheme, the biomass was left to settle for 60 min and then 1.35 L of supernatant was discarded. Thereafter, the reactor was filled with the same volume of soybean wastewater to start a new cycle. Operation consisted of one cycle per day. This feeding regime of every 1 d resulted in the following average hydraulic residence times (HRTs) for the four bioreactors run: 3 d (IF1), 3 d (IF2), 3 d (IF3) and 3 d (IF4) (see e-Supplementary Materials). The pH ranged from 4.5 to 8.5 and the DO level was controlled between 0.2 and 0.5 mg/L during the aerobic phase, which also helped minimize foaming. Each of the SBRs employed in this study was equipped with a magnetic stir plate to ensure mixed liquor homogeneity, a pair of EasySense pH and DO probes with their corresponding transmitters (Mettler Toledo), dedicated air and feed pumps, a solenoid valve for supernatant discharge, and a surrounding water jacket connected to a re-circulating water heater. The different portions of the cycle were controlled by a computer software specifically designed for these reactors (VentureMerger, Singapore). Water chemical analyses were done as described in Santillan et al. (2021).

### 2.3 Biomass protein and microbial analysis

The protein content of the biomass was quantified based on amino acid analysis using high-performance liquid chromatography (HPLC), following the procedure described by Vethathirri et al. (2023). Microbial community composition was assessed through 16S rRNA gene metabarcoding, as outlined in Vethathirri et al. (2023). For a total of 83 samples, an average of 49,108 reads were retained per sample after processing, constituting 66% of the average input reads. The adequacy of sequencing depth after reads processing was corroborated with rarefaction curves at the ASV level (see e-Supplementary Materials, Figure S1). Microbial characterization was further supported with fluorescence in situ hybridization (FISH) for domain bacteria and selected core SCP taxa (see e-Supplementary Materials).

### 2.4 Microbial protein production rate, yield, and nutrient removal efficiency

The biomass yield was calculated as the ratio of total suspended solids production and the total amount of soluble chemical oxygen demand (sCOD) or soluble total Kjeldahl nitrogen (sTKN) consumed through the feeding of wastewaters as given by Eq. (1) and Eq. (2), where TSS_n_ and TSS_0_ represent the final and initial mass of dry biomass, respectively. Similarly, the protein yield was calculated as the ratio of total protein production and the total amount of sCOD or sTKN consumed through the feeding of wastewaters as given by Eq. (3) and Eq. (4), where SCP_n_ and SCP_0_ represent the final and initial mass of biomass protein, respectively. The biomass production rate (Eq. (5)) was calculated as the ratio of total suspended solids production and the reactor working volume (L_R_) over the entire bioconversion period, *n* (in days). Likewise, the protein production rate (Eq. (6)) was calculated as the ratio of total protein production and the reactor working volume over the bioconversion period. Further, average removal efficiencies of nitrogen (R_sTN_) and carbon (R_sCOD_) present in the wastewaters were estimated as given by Eq. (7) and Eq. (8), respectively, where *m* represents the number of wastewater batches used for a reactor under the feeding scheme of this study, and sCOD_effluent,*m*_ and sTN_effluent,*m*,_ respectively, refer to the sCOD and sTN available in the reactor before the subsequent feeding when the *m^th^* wastewater batch was used in the previous feeding.

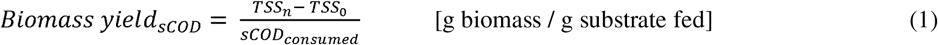

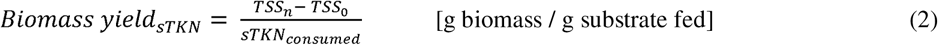

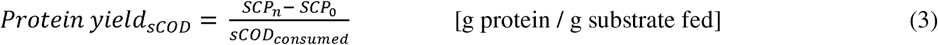

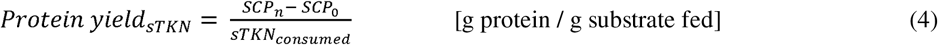

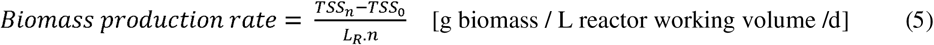

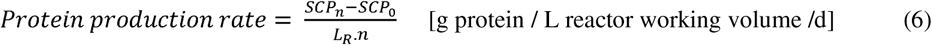

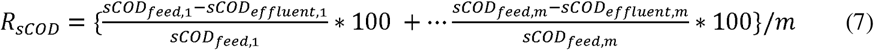

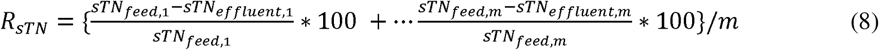

### 2.5 Statistical analyses and data visualization

Univariate tests through the two-tailed Welch’s t-test assuming unequal variances were done using the *rstatix* (v.0.6.0) R-package (Kassambara, 2020). All reported P-values for statistical tests in this study were corrected for multiple comparisons using a false-discovery rate (FDR) of 5% (Benjamini & Hochberg, 1995). Heat maps for bacterial relative abundances of the SCP were constructed using the DivComAnalyses package (v.0.9) in R (Constancias & Sneha-Sundar, 2022). Biomass growth trends were visualized using locally estimated scatterplot smoothing (*loess*) regression to describe overall temporal patterns in total suspended solids (TSS) and protein content. Plotting was performed in R (version 4.4.1) with the *ggplot2* and *ggpubr* packages. Individual data points for each reactor (IF1–IF4) were displayed with distinct symbols, and a single *loess* curve with 95% confidence intervals was fitted across all replicates to highlight the mean trajectory of biomass and protein accumulation over the 44-day operation period.

### 2.6 Optimization of heat treatment conditions for RNA reduction in SCP biomass

The heat treatment conditions for RNA reduction in single-cell protein were optimized by systematically evaluating three key factors: pH, aqueous matrix type, and temperature. The experiment was divided into four phases. In Phase 1, the effect of pH on RNA reduction and amino acid enrichment during heat treatment was evaluated. Dry SCP biomass samples were suspended in Milli-Q® water (initial pH of 9.7, measured using a calibrated pH meter). Acidic and alkaline conditions were established using volumetric dilutions of 0.1 M stock solutions of HCl and NaOH, respectively. The indicated percentages refer strictly to these volumetric dilutions (v/v) and do not represent arbitrary concentrations or mass fractions. For example, 50% HCl corresponds to a 1:1 (v/v) dilution of 0.1 M HCl with Milli-Q® water, while 1% NaOH corresponds to a 1:10 (v/v) dilution of 0.1 M NaOH prior to pH adjustment. The suspensions were adjusted to target pH values of 2.2 (100% HCl), 4.7 (50% HCl), 6.7 (40% HCl), and 9.8 (1% NaOH), with final values measured and confirmed using a calibrated pH meter rather than inferred from dilution ratios. In a separate condition, sodium chloride was added as an undiluted stock solution to modify ionic strength. This treatment did not serve as a pH-adjusting reagent; the suspension was already alkaline (initial pH 9.7), and the measured pH following NaCl addition was 9.3, reflecting minor ionic strength effects rather than acid–base neutralization. In Phase 2, the heat treatment at pH 10 in Milli-Q® water without additional chemical adjustment was conducted in three replicate samples. In Phase 3, different aqueous matrices (bioreactor effluent, Milli-Q® water, and tap water) were tested for suspending the dry bacterial SCP at 65 °C. In Phase 4, heat treatment temperatures ranging from 60 to 70 °C were applied.

Bioreactor effluent was collected from a 100-L SCP-producing SBR operated at room temperature under intermittent aeration (see e-Supplementary Materials, Table S2). The bioreactor was seeded with 20LL of microbial community-based inoculum generated from 4LL SBRs operated as described in Section 2.2. Bioreactor effluents containing bacterial biomass in the form of flocs were collected during the settling phase and centrifuged at 11,180Lg at 4 °C for 10 min. The supernatant was filtered using a 0.2 µm Acrodisc 25Lmm syringe filter (Pall Corporation) to remove residual biomass before SCP heat treatment. In all phases, biomass was first freeze-dried using a lyophilizer (Labconco) to constant weight; next, 0.5 g of dried biomass was suspended in 9.5 mL of aqueous matrix in a falcon tube and vortexed to obtain a 5% (w/v) homogenous biomass suspension. Further, the suspension was heated at the indicated test temperature in the water bath for 10 min followed by continuous incubation at 55° C for 2 h. The treated samples were then centrifuged at 11,180 g at 4° C for 10 min. The supernatant was decanted, and the pelleted biomass was subjected to freeze-drying to obtain dry biomass samples for subsequent quantification of nucleic acids and amino acids. The sample weights were recorded both prior to and following heat treatment. RNA from the heat-treated SCP samples were extracted and purified with a magnetic bead-based extraction method using the ZymoBIOMICS™ MagBead RNA kit (Cat # R2135) following the manufacturer’s instruction. Nucleic acid purity was determined as the absorbance ratio at wavelengths 260/280 using a NanoDrop 2000 UV-spectrophotometer (Thermo Scientific, USA). The concentrations of the purified samples were also measured with the Invitrogen Qubit™ 2.0 fluorometer using the Qubit™ RNA HS Assay Kit (Cat # Q32852) according to the manufacturer’s protocol. The total amino acids in the heat-treated SCP were analysed via HPLC after hydrolysis, identifying 21 amino acids through acid and base hydrolysis (Vethathirri et al., 2023).

## 3. Results and Discussion

### 3.1 Chemical composition and microbial characterization of influent wastewater

The soybean-processing wastewater used in this study was rich in organic carbon and contained variable concentrations of key nutrients, providing a suitable substrate for microbial protein production. The wastewater exhibited low alkalinity (0–131 mg LL¹) and an acidic pH (3.8–4.2), conditions favourable for heterotrophic rather than nitrifying activity (see e-Supplementary Materials, Table S1). Soluble chemical oxygen demand (sCOD) and total Kjeldahl nitrogen (sTKN) ranged from 2575 to 11,370 mg LL¹ and 21 to 109 mg LL¹, respectively, resulting in high C:N ratios (92–138) well above the recommended 10–20 range for balanced microbial growth (Vethathirri et al., 2021). Suspended solids were within 120–500 mg LL¹, and the native microbial community was dominated by lactic acid bacteria, mainly *Lactococcus*, *Weissella*, and *Lactobacillus* (see e-Supplementary Materials, Figure S2). These taxa are characteristic of soybean fermentation environments and represent microorganisms that may contribute to community-based SCP production. Furthermore, changes in community structure in soy-derived fermentation systems have been shown to enhance biosynthesis pathways of essential amino acids (e.g., valine, leucine, isoleucine) in later stages of fermentation (Meng et al., 2024). Given that lactic acid bacteria possess complete pathways for branched-chain amino acid biosynthesis, their dominance indicates a community metabolically oriented toward amino acid production, consistent with the observed protein accumulation (Figure 2).

**Figure 2.**
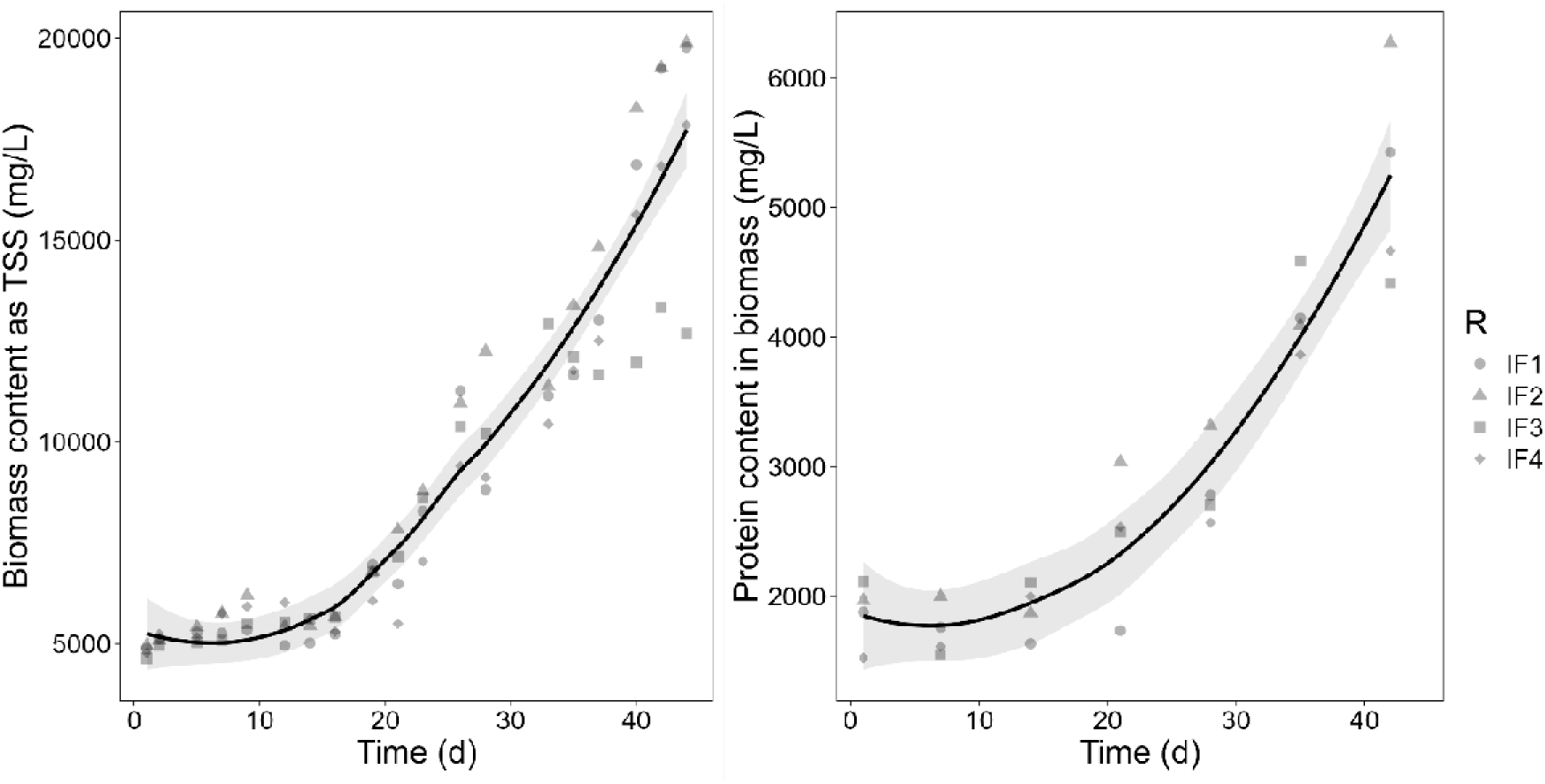
Biomass and protein accumulation in four replicate inoculated sequencing batch reactors (IF1–IF4) run for 44 days. The left panel shows biomass concentration expressed as total suspended solids (TSS, mg LLJ¹), and the right panel shows protein content in biomass (mg LLJ¹) quantified from amino acid analysis by HPLC. Each point represents one measurement per reactor, with distinct symbols indicating the four replicates. Smooth black lines show *loess* regression fits with 95% confidence intervals (grey shading). All reactors were seeded with the same enriched microbial community (initial TSS = 4815 mg LLJ¹) and run at a hydraulic retention time of 3 days.

### 3.2 Biomass growth and protein accumulation in microbial community-based SCP production

Sustained biomass growth and protein accumulation were achieved in all reactors throughout the 44-day operational period (Figure 2). Total suspended solids (TSS) increased steadily up to a maximum concentration of 14,920 mg LL¹, indicating efficient conversion of soybean-processing wastewater into microbial biomass. The four 4-L reactors were operated without biomass withdrawal to maximize yield, starting from an initial TSS of 4,815 mg LL¹ and an average food-to-microorganism ratio (F:M) of 0.25 g sCOD gL¹ TSS dL¹. Although the feed wastewater nitrogen content varied (sTKN = 21–109 mg LL¹) and C:N ratios were high (Table 1), no decline in biomass concentration was observed; this coincided with daily substrate input, which maintained continuous nutrient availability despite low nitrogen content in the influent.

**Table 1.**
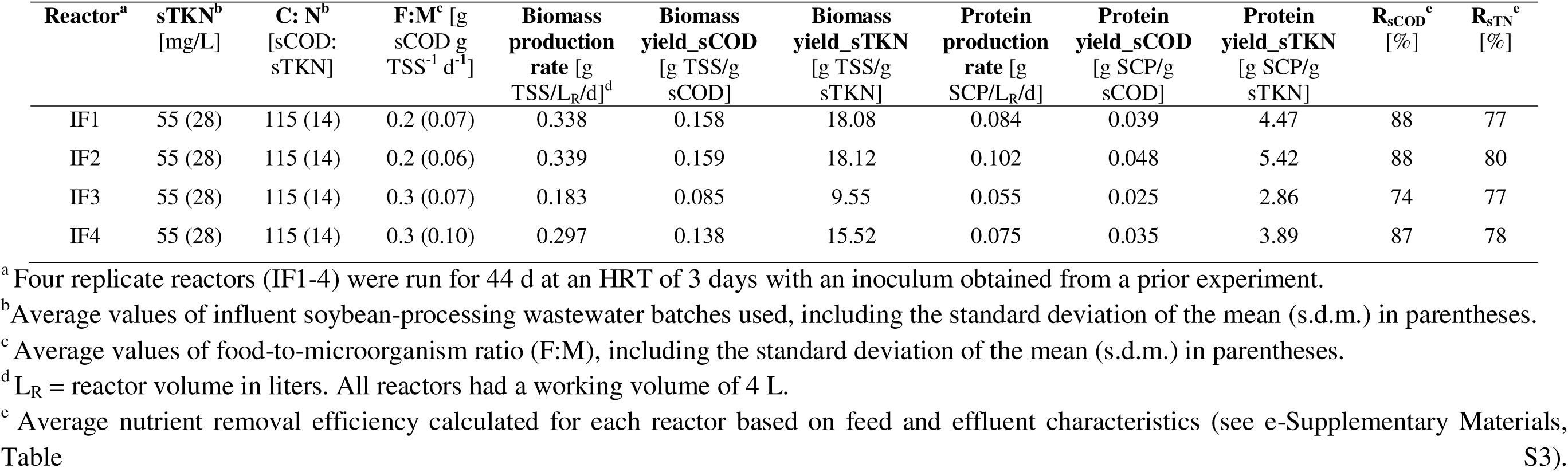
Single cell protein production from soybean-processing wastewaters in replicate reactors using a microbial community-based inoculum.

Protein content followed the same temporal pattern as biomass growth (Figure 2), increasing after day 19 when wastewater batches containing more nitrogen (61–109 mg LL¹ sTKN) were introduced. The microbial biomass comprised essential amino acids (EAA) required for aquaculture species (Salamanca et al., 2025) and reached a protein content 36 ± 1.8% on day 42 (see e-Supplementary Materials, Figure S3). Arginine was the most abundant EAA (10.2 ± 0.6% protein); it plays a key role in mitigating stress, supporting immune function, and enhancing growth in aquaculture species (Chuphal et al., 2025). Recent aquaculture trials with microbial community-based SCP (Santillan et al., 2024; Santillan et al., 2025; Santillan et al., 2026) indicate that the SCP produced in this study could serve as a sustainable feed ingredient, supplying essential amino acids for juvenile fish and shrimp and enhancing aquaculture resilience. Overall, reactors inoculated with an enriched microbial community and fed daily achieved consistent biomass and protein accumulation, demonstrating robust community-based SCP production even when influent nitrogen concentrations were low.

### 3.3 Impact of microbial inoculum and HRT on SCP production and effluent quality

Inoculation with an enriched microbial community and operation at a shorter hydraulic retention time (HRT) markedly improved SCP production without compromising effluent quality. The four inoculated bioreactors, seeded with biomass from a previous experiment using the same soybean-processing wastewater, achieved an average biomass production rate that was 426% higher than that if uninoculated reactors operated at HRT = 7.2 days (Vethathirri et al., 2025). The corresponding protein production rate increased by 251%, with reactor IF2 reaching the highest values of 0.339 g TSS LL¹ dL¹ and 0.102 g protein LL¹ dL¹ (Table 1). Biomass yields relative to sCOD and sTKN increased by 214% and 217%, respectively (Table 2). Similar improvements in productivity at shorter feeding intervals have been observed in other systems, such as mixed microalgae cultures (Solmaz & Işık, 2020). Despite the higher production rates, effluent treatment remained stable, with mean removal efficiencies of 84% sCOD and 78% sTN, comparable to uninoculated operation at a long HRT. These differences in yield and production rate were statistically significant (two-tailed Welch’s t-tests, p < 0.05) (Table 2), confirming that the combined effect of inoculation and more frequent feeding enhanced productivity while maintaining effluent quality. Although inoculated reactors at shorter HRT showed significantly higher biomass and protein production compared with uninoculated reactors at longer HRT, this effect was inferred from observed trends rather than directly tested. While the wastewater batches used in the different runs originated from the same soybean-processing facility and are therefore reasonably comparable, temporal variations in feedstock composition cannot be fully excluded as a contributing factor. Consequently, the observed trends are consistent with enhanced productivity due to inoculation and operational control, but the combined effect cannot be definitively confirmed without a dedicated control experiment designed to isolate these variables.

**Table 2.**
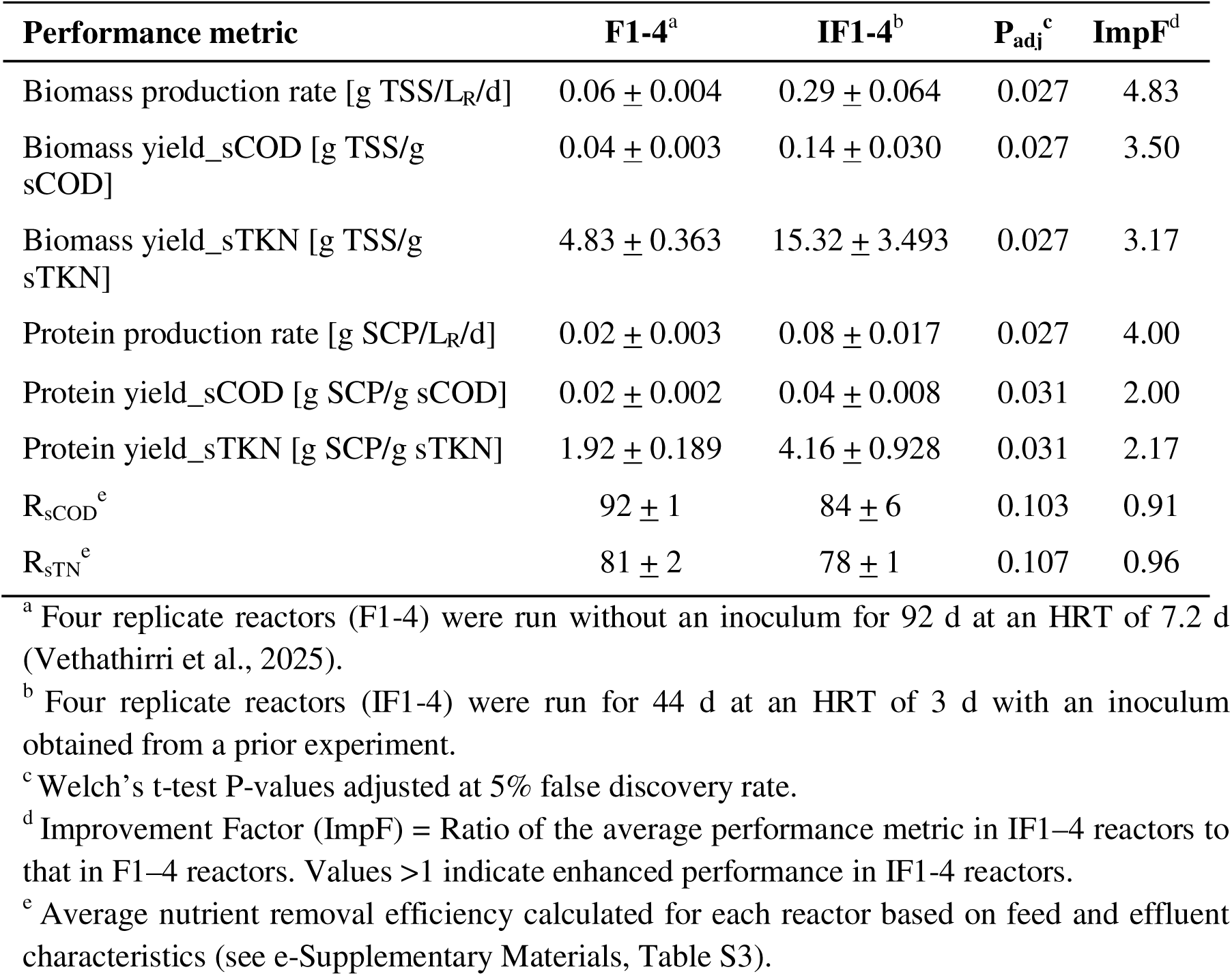
Comparison of the average performance of reactors run with and without inoculum in microbial community-based SCP production from soybean-processing wastewater. The HRT was 7.2 d in uninoculated reactors (F1-4) and 3 d in inoculated reactors (IF1-4) used in this study (see Table 1).

| Performance metric | F1-4<sup>a</sup> | IF1-4<sup>b</sup> | P<sub>adj</sub><sup>c</sup> | ImpF<sup>d</sup> |
| --- | --- | --- | --- | --- |
| Biomass production rate [g TSS/L<sub>R</sub>/d] | 0.06 ± 0.004 | 0.29 ± 0.064 | 0.027 | 4.83 |
| Biomass yield_sCOD [g TSS/g sCOD] | 0.04 ± 0.003 | 0.14 ± 0.030 | 0.027 | 3.50 |
| Biomass yield_sTKN [g TSS/g sTKN] | 4.83 ± 0.363 | 15.32 ± 3.493 | 0.027 | 3.17 |
| Protein production rate [g SCP/L<sub>R</sub>/d] | 0.02 ± 0.003 | 0.08 ± 0.017 | 0.027 | 4.00 |
| Protein yield_sCOD [g SCP/g sCOD] | 0.02 ± 0.002 | 0.04 ± 0.008 | 0.031 | 2.00 |
| Protein yield_sTKN [g SCP/g sTKN] | 1.92 ± 0.189 | 4.16 ± 0.928 | 0.031 | 2.17 |
| R<sub>sCOD</sub><sup>e</sup> | 92 ± 1 | 84 ± 6 | 0.103 | 0.91 |
| R<sub>sTN</sub><sup>e</sup> | 81 ± 2 | 78 ± 1 | 0.107 | 0.96 |

<sup>a</sup> Four replicate reactors (F1-4) were run without an inoculum for 92 d at an HRT of 7.2 d (Vethathirri et al., 2025).
<sup>b</sup> Four replicate reactors (IF1-4) were run for 44 d at an HRT of 3 d with an inoculum obtained from a prior experiment.
<sup>c</sup> Welch's t-test P-values adjusted at 5% false discovery rate.
<sup>d</sup> Improvement Factor (ImpF) = Ratio of the average performance metric in IF1-4 reactors to that in F1-4 reactors. Values >1 indicate enhanced performance in IF1-4 reactors.
<sup>e</sup> Average nutrient removal efficiency calculated for each reactor based on feed and effluent characteristics (see e-Supplementary Materials, Table S3).

### 3.4 Microbial community shifts and dominant taxa under inoculated SCP operation

The most abundant SCP-producing genera in reactors included *Acidipropionibacterium*, *Lactococcus*, *Megasphaera*, *Lactobacillus*, and *Azospirillum* along with ASVs identified at higher taxonomic levels, including *f_Weeksellaceae* and *o_Saccharimonadales* (Figure 3a). Notably, the *f_Weeksellaceae* ASV was abundant across all reactors, while the *o_Saccharimonadales* ASV appeared among the top 30 in specific reactors, suggesting these taxa represent core community members under reactor conditions, although their specific functional roles remain uncharacterized at present. Fluorescence in situ hybridization (FISH) imaging confirmed the presence of *Acidipropionibacterium* and *Lactococcus* within the SCP biomass (Figure 3b). *Acidipropionibacterium* and *Azospirillum* were absent among the top 30 genera in the influent soybean-processing wastewater (see e-Supplementary Materials, Figure S2), yet *Acidipropionibacterium* and *Megasphaera*, dominant in the initial inoculum, became core members of the reactor communities (up to 9.2% and 12.5% relative abundance, respectively). Similar inoculum-derived dominance has been reported in other community-based protein production systems (Hülsen et al., 2018a; Hülsen et al., 2018b; Hülsen et al., 2020). The low dissolved oxygen levels (0.2–0.5 mg LL¹) selectively favored fermentative and microaerophilic taxa such as *Megasphaera*, a vitamin B_12_ producer (Balabanova et al., 2021; Nallabelli et al., 2016), while under nitrogen-limited conditions, taxa capable of efficient nitrogen assimilation or associative nitrogen fixation, such as *Azospirillum*, gain a competitive advantage (Vethathirri et al., 2023). These findings underscore that nutrient availability and reactor conditions can play a more critical role in shaping microbial community composition during SCP production than bacterial immigration from the influent wastewater (Vethathirri et al., 2023; Vethathirri et al., 2025).

**Figure 3.**
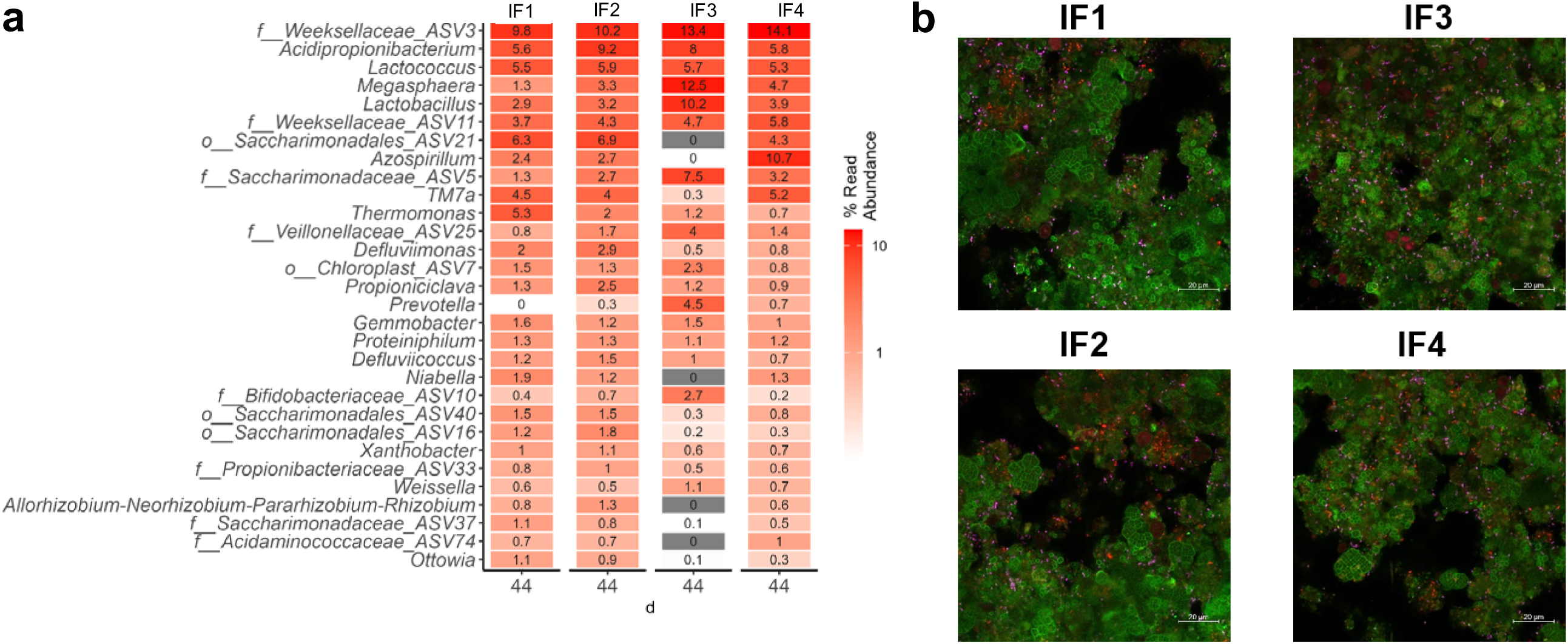
Microbial characterization of community-based SCP biomass produced from soybean-processing wastewater. (**a**) The 30 most abundant bacterial genera and ASVs in four replicate reactors (IF1-4) that were run for 44 d and received an inoculum based on 16S rRNA gene amplicon sequencing. Data correspond to samples collected on d44. Note that ASVs identified at the family (f_) or order (o_) level, which had sufficient abundance to be ranked among the top 30, are included alongside genera. (**b**) FISH images from four replicate reactors (IF1-4) that received an inoculum on d42 showing *Lactococcus lactis* (magenta), *Acidipropionibacterium* (red) and all bacterial (green) cells. Cell abundances are consistent with 16S rRNA gene amplicon sequencing data, showing on average 10.2% relative abundance for *Lactococcus lactis*. Similarly, for *Acidipropionibacterium*, cell counts are consistent with 16S rRNA gene amplicon sequencing data, showing on average 12.0% relative abundance. Additional reactor samples were analyzed for high temporal resolution throughout the study (see e-Supplementary Materials, Figure S4).

### 3.5 Effluent-assisted RNA reduction and nutritional refinement of microbial protein

Heat treatment effectively reduced RNA content in microbial community-based protein produced from soybean-processing wastewater, with outcomes varying based on pH, aqueous matrix type and temperature (Figure 4). However, RNA reduction is not universally required in aquaculture applications, where moderate levels can enhance immunity and growth (Burrells et al., 2001; Li & Gatlin., 2006). High nucleic acid concentrations are more relevant in human or pet food contexts, where excess purine intake may elevate uric acid or overestimate true protein content (Hadi & Brightwell., 2021; Zhuang et al., 2024). Thus, the refinement step evaluated here is most applicable when lower RNA levels are desired, and its implementation should be guided by end-use considerations.

**Figure 4.**
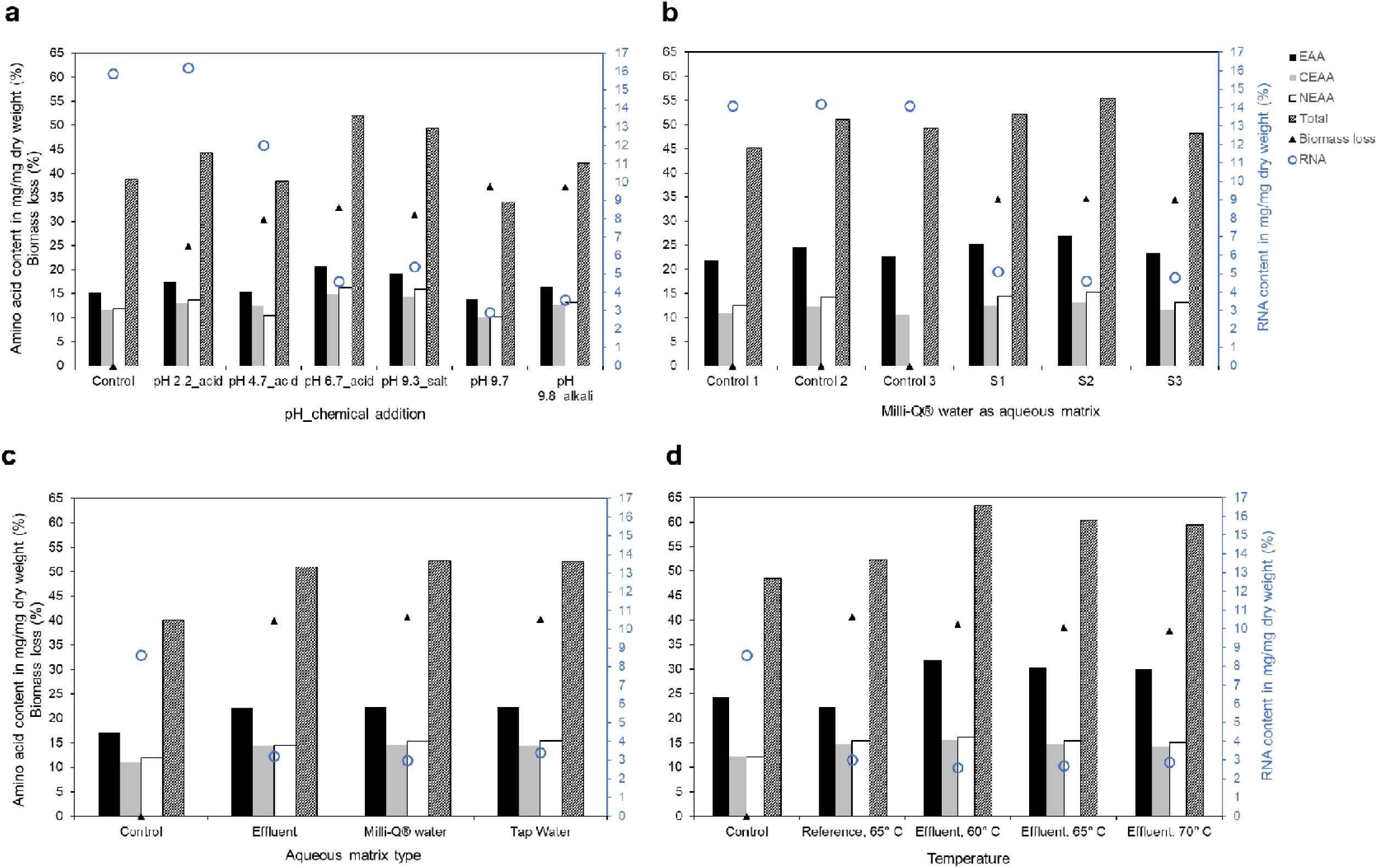
Amino acid (bars, left y-axis), biomass loss (solid triangles, left y-axis), and RNA (open blue circles, right y-axis) contents in microbial community-based protein produced from soybean-processing wastewater using inoculated consortia. **(a)** Comparison between control and heat-treated samples under different pH conditions: pH 2.2, 4.7, 6.7, 9.8, and initial suspension in Milli-Q® water (pH 9.7). **(b)** Comparison of control (n = 3) and heat-treated replicate samples (n = 3; S1-S3) using Milli-Q® water without pH adjustment. (**c**) Comparison between control and heat-treated samples using different aqueous matrix types: bioreactor effluent, Milli-Q® water, and tap water. (**d**) Comparison among control, reference and bioreactor effluent-treated samples subjected to different heat treatment temperatures: 60, 65, and 70LJ°C. Reference denotes SCP samples treated with Milli-Q® water at 65 °C in the same trial run. Controls in panels (a) to (d) refer to the freeze-dried sludge sample directly harvested from the reactor without any post-treatment and are included as baseline samples rather than test conditions for aqueous matrix and temperature comparisons, respectively. Biomass loss resulting from heat treatment was quantified and annotated for each test condition to represent the overall loss corresponding to each treatment condition and to illustrate trade-offs between RNA reduction and biomass recovery. Amino acid profiles are grouped as follows: EAA, essential amino acids (solid black square); CEAA, conditionally essential amino acids (solid grey square); NEAA, non-essential amino acids (open white square); TAA, total amino acids (patter-filled black square). Amino acid classifications are based on their essentiality for fish (Salamanca, 2025).

Among all pH conditions tested, the highest RNA reduction (∼82%) was achieved in the sample suspended in Milli-Q® water at pH 9.7 without the addition of any chemical solution (Figure 4a). A replicative analysis of the heat treatment in Milli-Q® water showed a consistent RNA reduction of 65.8 ± 1.5% and an average increase in total amino acids of 7.1 ± 6.2% (Figure 4b). At 65 °C, samples treated with Milli-Q® water, bioreactor effluent, and tap water achieved RNA reductions of 65%, 63%, and 60%, respectively (Figure 4c). All treatments increased amino acid content, with enrichments of 30%, 27%, and 30%, while biomass loss was lowest in the effluent-treated sample (39.9%) compared to 40.3% for tap water and 40.7% for Milli-Q® water. While Milli-Q® water yielded a slightly higher RNA reduction and amino acid enrichment, bioreactor effluent offered a more sustainable trade-off by minimizing biomass loss without substantially compromising SCP quality.

When temperature was varied using the effluent as aqueous matrix, RNA decreased by 70%, 69%, and 66% at 60, 65, and 70 °C, respectively, while the reference treated with Milli-Q® water at 65 °C showed a lower reduction of 65% (Figure 4d). Amino acid enrichment across effluent-treated samples ranged from 22% to 30%, with consistent biomass losses of about 39%, all with total amino acid contents in the range of 59–63% and no observable negative effect on essential amino acids relative to other fractions. Overall, the RNA levels observed in the untreated control samples (12.6 ± 2.9%) are consistent with the high nucleic acid contents typically reported for bacterial SCP (8–12%) (Ritala et al., 2017), which often necessitate processing prior to use in food or feed applications (Felix et al., 2023; Santillan et al., 2025). When bioreactor effluent was used as the aqueous matrix for heat treatment at 60 °C, the RNA content was reduced from 8.6% to 2.6%, accompanied by a 30% increase in total amino acids despite a biomass loss of 39%, demonstrating the effectiveness of effluent-assisted refinement.

Complete RNA removal may even be undesirable, as moderate dietary RNA can act as an immunostimulant, improving fish and shrimp disease resistance and survival under stress (Rairat et al., 2022; Alloul et al., 2021). Beyond its nutritional role, RNA may also contribute to the postbiotic potential of microbial community-based SCP, as microbial nucleic acids, cell wall fragments, and metabolites can modulate gut immunity and host responses (Wegh et al., 2019; Meena et al., 2025). The comparatively lower RNA reduction observed in this study, particularly with bioreactor effluent, likely reflects the presence of salts and organics that stabilize RNases and slow thermal denaturation (Baba et al., 2017), but it also aligns with a sustainability-driven approach where mild treatment preserves product bioactivity. Integrating such optional refinement within the same process framework broadens the applicability of community-based SCP to diverse feed and food contexts. Reusing internal effluent streams for downstream product improvement reduces reliance on fresh water and chemical additives while maintaining product quality, reinforcing the dual-loop framework as a resource-efficient and functional bioprocess for sustainable protein production.

The thermal refinement step (60 °C) effectively reduced RNA content and increased total amino acid concentration; however, it introduces an additional energy demand that must be considered in process scale-up. Although the operating temperature is moderate compared to heat sterilization, energy consumption and heat transfer efficiency will influence overall process sustainability. In industrial settings, this step could potentially be integrated with low-grade or residual heat streams from adjacent operations, thereby reducing net energy input. Heat recovery systems or continuous-flow heat exchanger configurations may further improve thermal efficiency. A quantitative energy balance and techno-economic assessment were beyond the scope of the study but will be necessary to evaluate the cost-benefit ratio and optimize the sustainability of the refinement loop within the integrated dual-loop framework.

### 3.6 Future directions for optimizing community-based SCP production from food-processing wastewater

This study employed an HRT of 3 days and inoculation with an enriched microbial community from the same soybean-processing wastewater source, resulting in marked improvements in yield and productivity compared with a longer HRT and the absence of inoculation (Vethathirri et al., 2025) (Table 2). Although SCP yield and production rate increased significantly, the study was not designed to test the individual effects of HRT and inoculation, which should be evaluated independently in future studies. Shorter HRTs may further enhance biomass recovery by improving settling capacity and increasing biomass concentration, as observed under high-rate activated sludge conditions at reduced HRTs (AlSayed et al., 2023). For instance, purple non-sulfur bacterial biomass produced in an anaerobic upflow photobioreactor aggregated efficiently at an HRT of 0.1 days but disaggregated at an HRT of 1 day, an effect attributed to unidentified heat-labile metabolites (Blansaer et al., 2022). Further, nitrogen limitation likely constrained biomass accumulation in this study, suggesting that adjustments in feeding pattern or substrate load are needed to sustain higher TSS concentrations while maintaining a balanced nutrient supply. Additionally, revising the feeding regime could help maintain the same food-to-microorganism ratio and support consistent reactor performance. Future research should explore combinations of reduced HRT and intermittent biomass harvesting, which have been shown to increase algal biomass productivity by more than 1.5-fold at an HRT of 2 days (Wang et al., 2022). Such optimization could maximize SCP yield and effluent quality while improving process stability, supporting the broader goal of developing resource-efficient and circular microbial protein bioprocesses from wastewaters produced in the food and beverage industry.

Although the dual-loop system demonstrated stable operation and performance in 4 L laboratory-scale reactors, scale-up will require careful engineering assessment, since this scale was selected to facilitate mechanistic evaluation and hypothesis testing. In Loop 1, increasing the reactor volume may affect mixing homogeneity, biomass retention, and microbial community stability, potentially influencing protein content of single-cell protein and nutrient removal efficiency. Where aerobic conditions are applied, oxygen transfer efficiency and solids separation capacity may impose additional design constraints. In Loop 2, heat transfer efficiency during the 60 °C biomass refinement step represents a critical scale-up consideration. Larger working volumes may introduce temperature gradients and increased energy demand, which could affect RNA reduction efficiency and product consistency. Implementation at larger scale would therefore require optimized reactor geometry, effective heat exchanger integration, and energy recovery strategies. Consequently, pilot-scale studies are necessary to validate process control, thermal management, and overall energy performance prior to industrial application.

## 4. Conclusions

This study introduced an integrated dual-loop bioprocess that couples microbial production, effluent reuse, and product refinement within a circular framework. Combining inoculum enrichment with moderate-cycle reactor operation enabled stable protein production from food-sector wastewaters while maintaining effective nutrient recovery. The enrichment patterns observed suggest that different microbial communities may offer comparable potential for achieving desirable protein yields and amino acid profiles under suitable operational conditions. Effluent-assisted heat treatment proved useful for reducing RNA levels and improving amino acid profiles, although at the expense of biomass yield and additional energy input. These results collectively underscore the trade-offs between process efficiency, nutritional quality, and resource use, and point toward the need for future innovations that improve energy efficiency and valorisation of internal process streams. The dual-loop framework thus provides a conceptual basis for designing sustainable bioprocesses for waste biomass management.

E-supplementary data for this work can be found in the e-version of this paper online.

## Data availability

DNA sequencing data are available at NCBI BioProjects PRJNA1372396. See supplementary information for details about HRT estimation, chemical characteristics of bioreactor effluent and influent, rarefaction plots for 16S rRNA gene sequencing data, essential amino acid profiles in reactors, and temporal dynamics of the 30 most abundant genera in each reactor.

