## Supplementary Information for "Dual-loop involving microbial single-cell protein production from soybean-processing wastewater and effluent-based refinement for circular bioeconomy applications"

Hydraulic retention time

Following the study design, after every 24 hours, the biomass was left to settle for 60 min, after which 1.35-L of the supernatant effluent was removed. The reactor was then filled with the same volume of soybean wastewater as feed. Reactors experienced different growth and sCOD removal rates due to the variable nutrient quality of the different soybean wastewater batches used to feed them. Additionally, as these reactors were designed for SCP accumulation, no biomass was removed from them during the study. Hence, an average hydraulic retention time (HRT) was estimated using Eq. (1):

$$HRT = \left\{\frac{V}{Q}\right\} \text{ [d]} \quad (1)$$

Where, V represents the working volume of the reactor (4 L), and Q refers to the flow rate of influent wastewater during each feeding episode (1.35 L/d).

Fluorescence in situ hybridization

Microbial characterization was further supported with fluorescence in situ hybridization (FISH) for domain bacteria and selected core SCP taxa. Samples were obtained from the reactor and fixed using 4% paraformaldehyde for 2 to 3 h on ice. The fixed samples were washed with 1× phosphate-buffered saline (PBS) solution and stored in a mixture of 1×PBS and ethanol (1:1) at -20 °C until use. The cells were allowed to dry on microscopic slides and dehydrate in an ethanol series of 50, 80 and 96% for 3 min each. Hybridization buffer (0.9 M NaCl, 20 mM Tris-HCl, 35% formamide, 0.01% sodium dodecyl sulfate) and probes were added to detect microorganisms of interest. Eubmix (Eub338, Eub 338ll, Eub 338lll) targeted most bacteria (Daims et al., 1999), AZOI 655 probe targeted species belonging to *Azospirillum* (Stoffels et al., 2001) and LactV5 probe targeted *Lactococcus lactis* (Ercolini et al., 2003). To target all species belonging to *Acidipropionibacterium*, probe 997 was obtained using ARB SILVA (Ludwig et al., 2004) and compared with the 16S rRNA sequences recovered from the reactors. After hybridization, the slides were washed with warm buffer for 10 min (0.9 M NaCl, 20 Mm Tris-HCl, 5 mM EDTA, 0.01% sodium dodecyl sulfate) and rinsed thoroughly with cold water. FISH images were acquired using an LSM780 confocal laser scanning microscope. The Zen software was used for image processing and graphical analysis (Carl Zeiss, German).

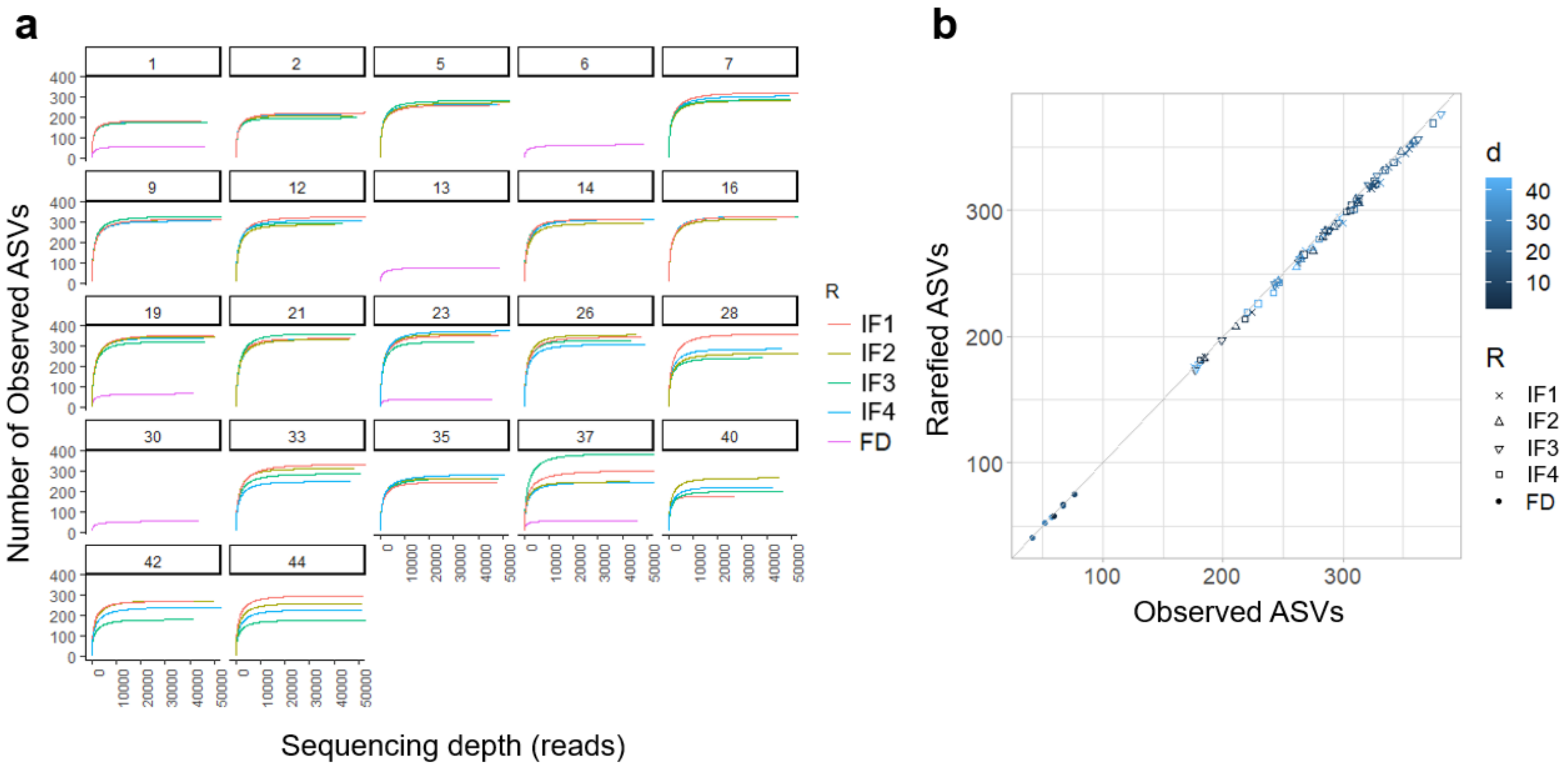


**Figure S1.** Rarefaction plots for 16S rRNA gene sequencing data. (**a**) Rarefaction curves for reactors and soybean-processing wastewater batches (FD) across time (**b**) Rarefied versus observed number of ASVs. Four reactors (IF1-4) were run for 44 d with inoculum under identical conditions.

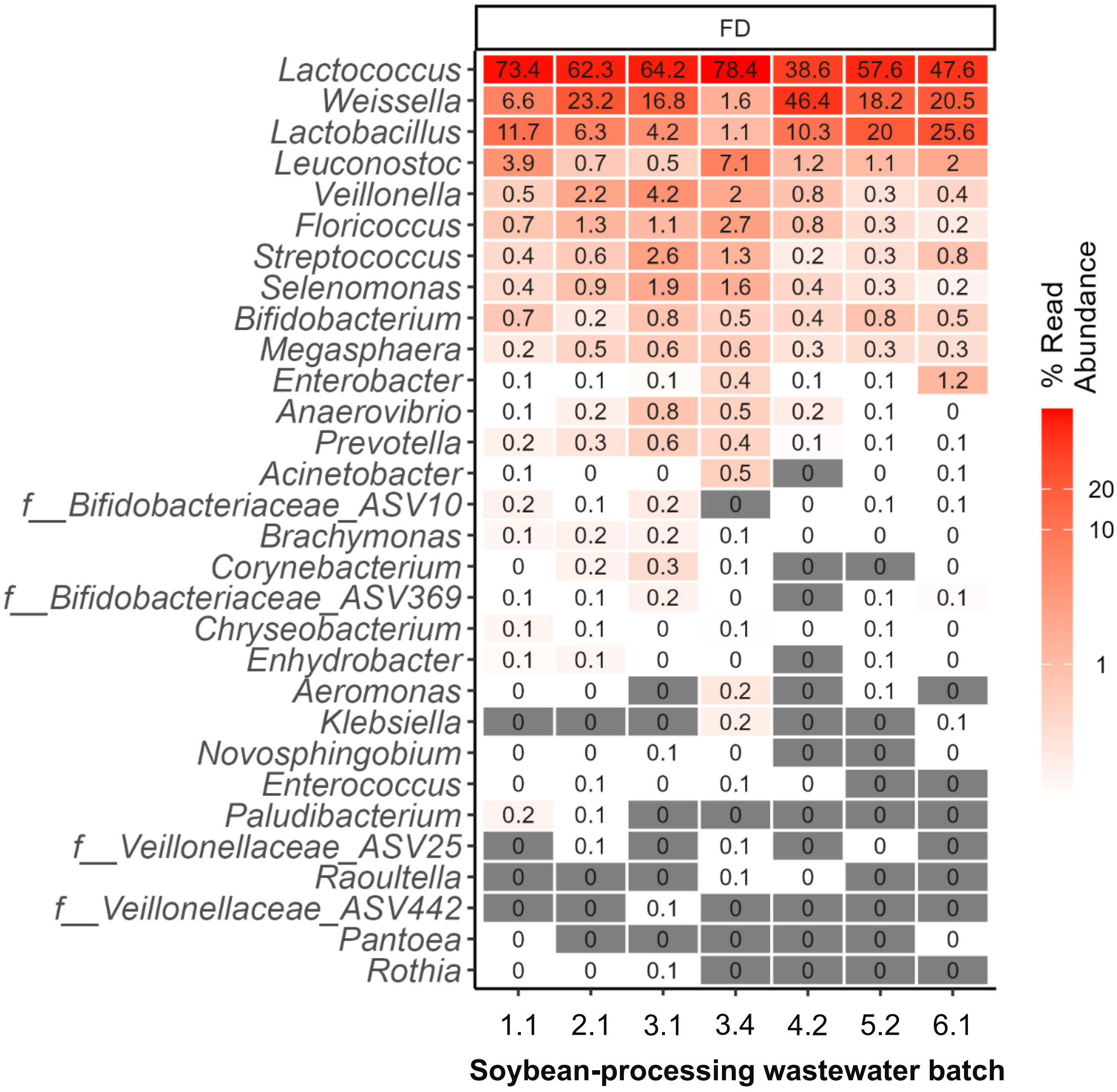


**Figure S2.** Microbial characterization of influent soybean-processing wastewaters used for microbial community-based SCP production, as assessed through 16S rRNA gene amplicon sequencing. Shown are the top 30 most abundant taxa, including genera and ASVs that could not be assigned at the genus level but were identified at the family (f_) level, for selected wastewater samples. All batches except Batch 3 consisted of two carboys, while Batch 3 consisted of four carboys. For the analysis, one representative carboy was selected from each batch, except for Batch 3, from which two carboys (3.1 and 3.4) were included. The samples shown are 1.1, 2.1, 3.1, 3.4, 4.2, 5.2, and 6.1 (see Table S1).

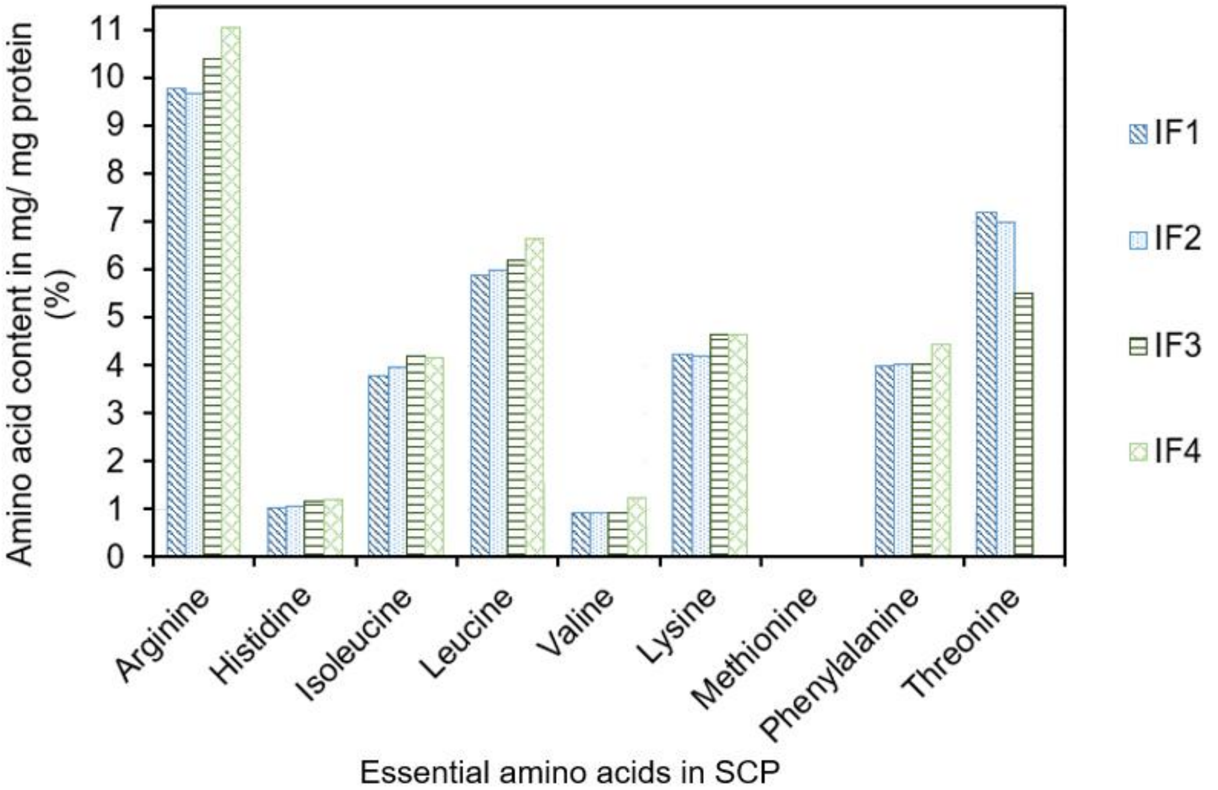


**Figure S3.** Essential amino acid composition of microbial community-based single-cell protein produced from soybean-processing wastewater for aquaculture applications. Four replicate reactors were run with inoculum: IF1 (pattern-filled dark blue bar), IF2 (pattern-filled light blue bar), IF3 (pattern-filled dark green bar), and IF4 (pattern-filled light green bar). Data used for all reactors correspond to samples collected on d42.

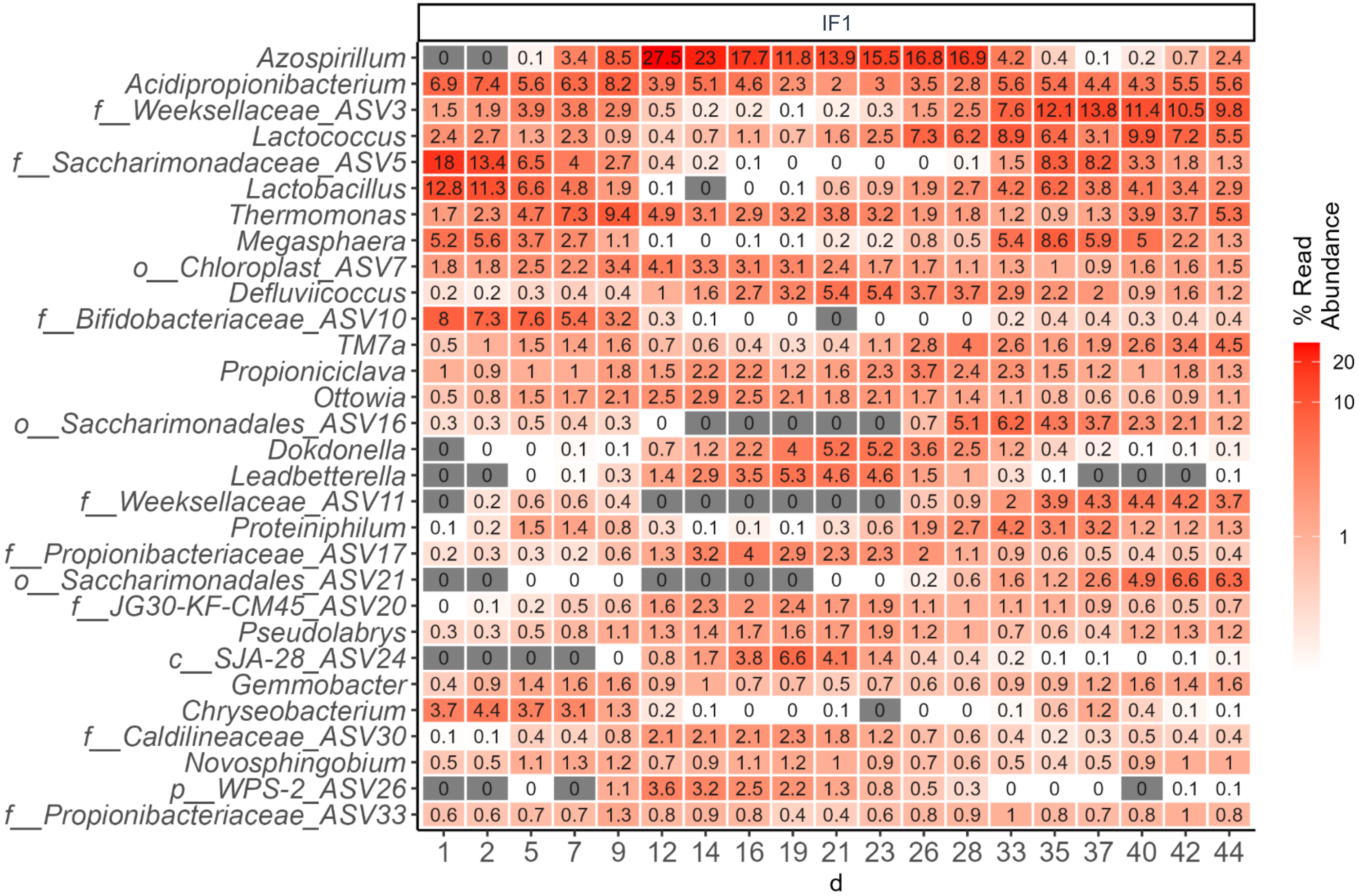

IF1
Azospirillum
Acidipropionibacterium
f__Weeksellaceae_ASV3
Lactococcus
f__Saccharimonadaceae_ASV5
Lactobacillus
Thermomonas
Megasphaera
o__Chloroplast_ASV7
Defluviicoccus
f__Bifidobacteriaceae_ASV10
TM7a
Propioniciclava
Ottowia
o__Saccharimonadales_ASV16
Dokdonella
Leadbetterella
f__Weeksellaceae_ASV11
Proteiniphilum
f__Propionibacteriaceae_ASV17
o__Saccharimonadales_ASV21
f__JG30-KF-CM45_ASV20
Pseudolabrys
c__SJA-28_ASV24
Gemmobacter
Chryseobacterium
f__Caldilineaceae_ASV30
Novosphingobium
p__WPS-2_ASV26
f__Propionibacteriaceae_ASV33
1 2 5 7 9 12 14 16 19 21 23 26 28 33 35 37 40 42 44
d
% Read Abundance
20
10
1

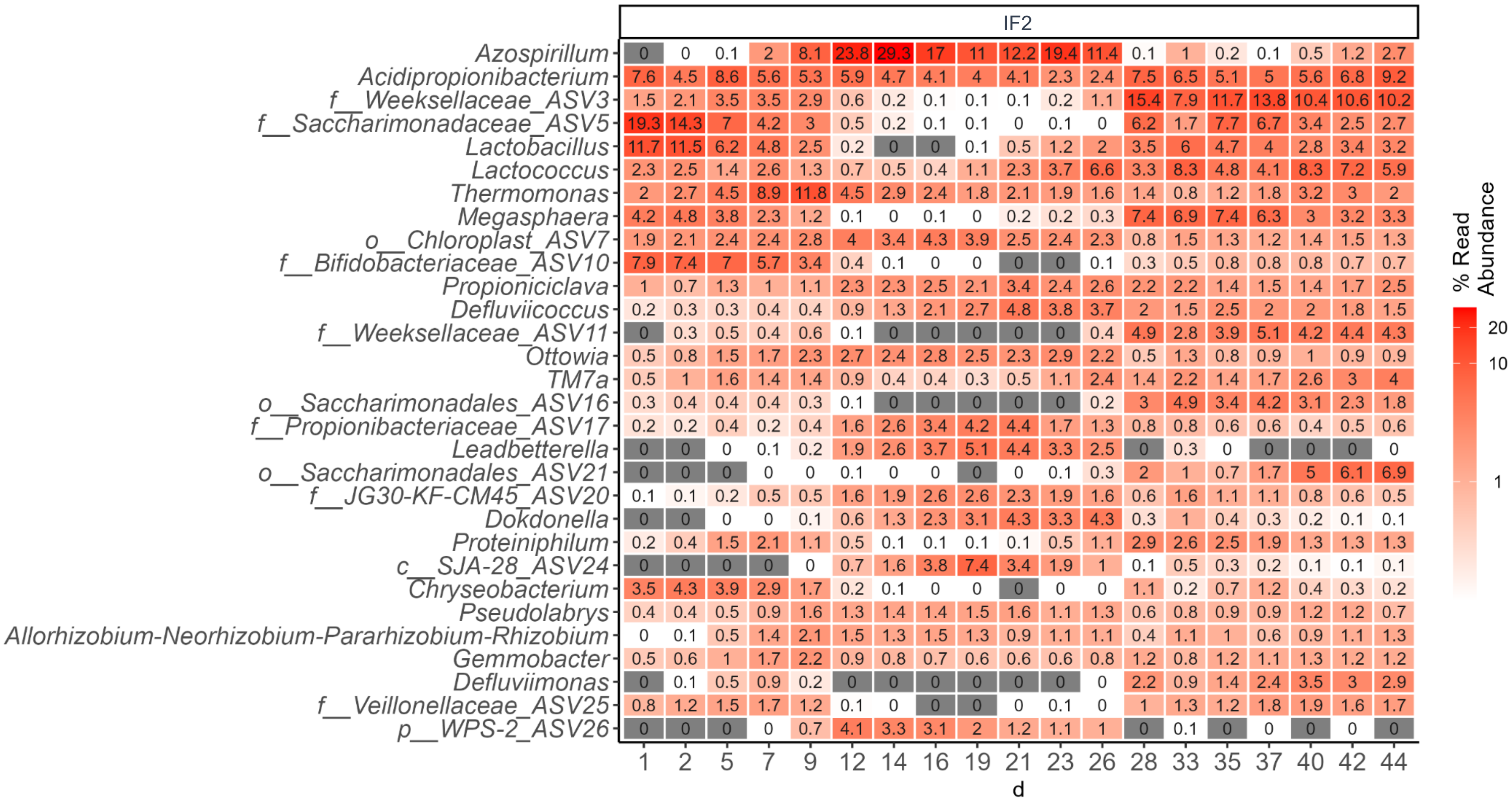
IF2
Azospirillum 0 0 0.1 2 8.1 23.8 29.3 17 11 12.2 19.4 11.4 0.1 1 0.2 0.1 0.5 1.2 2.7
Acidipropionibacterium 7.6 4.5 8.6 5.6 5.3 5.9 4.7 4.1 4 4.1 2.3 2.4 7.5 6.5 5.1 5 5.6 6.8 9.2
f__Weeksellaceae_ASV3 1.5 2.1 3.5 3.5 2.9 0.6 0.2 0.1 0.1 0.1 0.2 1.1 15.4 7.9 11.7 13.8 10.4 10.6 10.2
f__Saccharimonadaceae_ASV5 19.3 14.3 7 4.2 3 0.5 0.2 0.1 0.1 0 0.1 0 6.2 1.7 7.7 6.7 3.4 2.5 2.7
Lactobacillus 11.7 11.5 6.2 4.8 2.5 0.2 0 0 0.1 0.5 1.2 2 3.5 6 4.7 4 2.8 3.4 3.2
Lactococcus 2.3 2.5 1.4 2.6 1.3 0.7 0.5 0.4 1.1 2.3 3.7 6.6 3.3 8.3 4.8 4.1 8.3 7.2 5.9
Thermomonas 2 2.7 4.5 8.9 11.8 4.5 2.9 2.4 1.8 2.1 1.9 1.6 1.4 0.8 1.2 1.8 3.2 3 2
Megasphaera 4.2 4.8 3.8 2.3 1.2 0.1 0 0.1 0 0.2 0.2 0.3 7.4 6.9 7.4 6.3 3 3.2 3.3
o__Chloroplast_ASV7 1.9 2.1 2.4 2.4 2.8 4 3.4 4.3 3.9 2.5 2.4 2.3 0.8 1.5 1.3 1.2 1.4 1.5 1.3
f__Bifidobacteriaceae_ASV10 7.9 7.4 7 5.7 3.4 0.4 0.1 0 0 0 0 0.1 0.3 0.5 0.8 0.8 0.8 0.7 0.7
Propioniciclava 1 0.7 1.3 1 1.1 2.3 2.3 2.5 2.1 3.4 2.4 2.6 2.2 2.2 1.4 1.5 1.4 1.7 2.5
Defluviicoccus 0.2 0.3 0.3 0.4 0.4 0.9 1.3 2.1 2.7 4.8 3.8 3.7 2 1.5 2.5 2 2 1.8 1.5
f__Weeksellaceae_ASV11 0 0.3 0.5 0.4 0.6 0.1 0 0 0 0 0 0.4 4.9 2.8 3.9 5.1 4.2 4.4 4.3
Ottowia 0.5 0.8 1.5 1.7 2.3 2.7 2.4 2.8 2.5 2.3 2.9 2.2 0.5 1.3 0.8 0.9 1 0.9 0.9
TM7a 0.5 1 1.6 1.4 1.4 0.9 0.4 0.4 0.3 0.5 1.1 2.4 1.4 2.2 1.4 1.7 2.6 3 4
o__Saccharimonadales_ASV16 0.3 0.4 0.4 0.4 0.3 0.1 0 0 0 0 0 0.2 3 4.9 3.4 4.2 3.1 2.3 1.8
f__Propionibacteriaceae_ASV17 0.2 0.2 0.4 0.2 0.4 1.6 2.6 3.4 4.2 4.4 1.7 1.3 0.8 0.8 0.6 0.6 0.4 0.5 0.6
Leadbetterella 0 0 0 0.1 0.2 1.9 2.6 3.7 5.1 4.4 3.3 2.5 0 0.3 0 0 0 0 0
o__Saccharimonadales_ASV21 0 0 0 0 0 0.1 0 0 0 0 0.1 0.3 2 1 0.7 1.7 5 6.1 6.9
f__JG30-KF-CM45_ASV20 0.1 0.1 0.2 0.5 0.5 1.6 1.9 2.6 2.6 2.3 1.9 1.6 0.6 1.6 1.1 1.1 0.8 0.6 0.5
Dokdonella 0 0 0 0 0.1 0.6 1.3 2.3 3.1 4.3 3.3 4.3 0.3 1 0.4 0.3 0.2 0.1 0.1
Proteiniphilum 0.2 0.4 1.5 2.1 1.1 0.5 0.1 0.1 0.1 0.1 0.5 1.1 2.9 2.6 2.5 1.9 1.3 1.3 1.3
c__SJA-28_ASV24 0 0 0 0 0 0.7 1.6 3.8 7.4 3.4 1.9 1 0.1 0.5 0.3 0.2 0.1 0.1 0.1
Chryseobacterium 3.5 4.3 3.9 2.9 1.7 0.2 0.1 0 0 0 0 0 1.1 0.2 0.7 1.2 0.4 0.3 0.2
Pseudolabrys 0.4 0.4 0.5 0.9 1.6 1.3 1.4 1.4 1.5 1.6 1.1 1.3 0.6 0.8 0.9 0.9 1.2 1.2 0.7
Allorhizobium-Neorhizobium-Pararhizobium-Rhizobium 0 0.1 0.5 1.4 2.1 1.5 1.3 1.5 1.3 0.9 1.1 1.1 0.4 1.1 1 0.6 0.9 1.1 1.3
Gemmobacter 0.5 0.6 1 1.7 2.2 0.9 0.8 0.7 0.6 0.6 0.6 0.8 1.2 0.8 1.2 1.1 1.3 1.2 1.2
Defluviimonas 0 0.1 0.5 0.9 0.2 0 0 0 0 0 0 0 2.2 0.9 1.4 2.4 3.5 3 2.9
f__Veillonellaceae_ASV25 0.8 1.2 1.5 1.7 1.2 0.1 0 0 0 0 0.1 0 1 1.3 1.2 1.8 1.9 1.6 1.7
p__WPS-2_ASV26 0 0 0 0 0.7 4.1 3.3 3.1 2 1.2 1.1 1 0 0.1 0 0 0 0 0
1 2 5 7 9 12 14 16 19 21 23 26 28 33 35 37 40 42 44
d
% Read Abundance
20
10
1

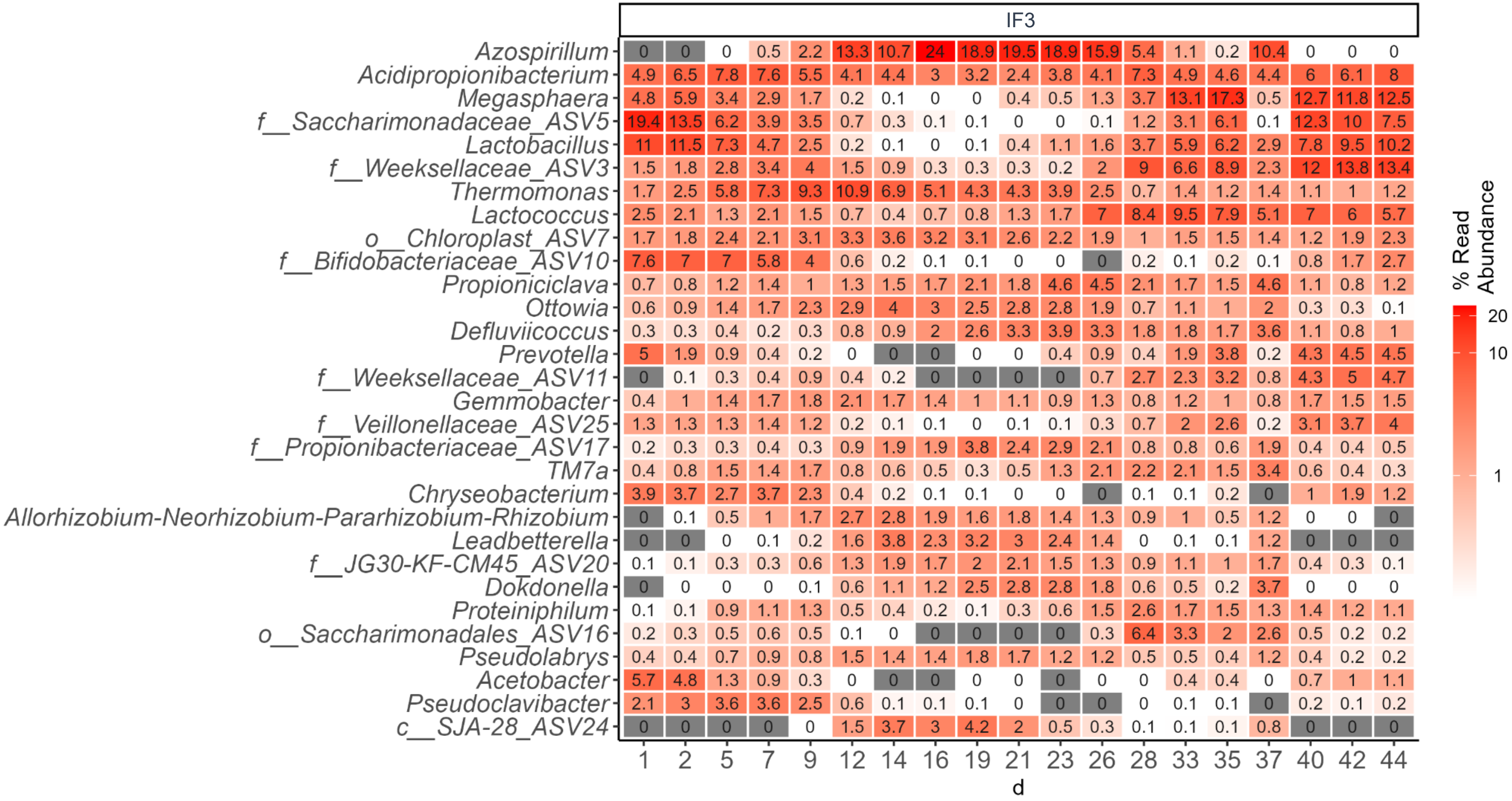
IF3
Azospirillum
Acidipropionibacterium
Megasphaera
f__Saccharimonadaceae_ASV5
Lactobacillus
f__Weeksellaceae_ASV3
Thermomonas
Lactococcus
o__Chloroplast_ASV7
f__Bifidobacteriaceae_ASV10
Propioniciclava
Ottowia
Defluviicoccus
Prevotella
f__Weeksellaceae_ASV11
Gemmobacter
f__Veillonellaceae_ASV25
f__Propionibacteriaceae_ASV17
TM7a
Chryseobacterium
Allorhizobium-Neorhizobium-Pararhizobium-Rhizobium
Leadbetterella
f__JG30-KF-CM45_ASV20
Dokdonella
Proteiniphilum
o__Saccharimonadales_ASV16
Pseudolabrys
Acetobacter
Pseudoclavibacter
c__SJA-28_ASV24
0 0 0 0.5 2.2 13.3 10.7 24 18.9 19.5 18.9 15.9 5.4 1.1 0.2 10.4 0 0 0
4.9 6.5 7.8 7.6 5.5 4.1 4.4 3 3.2 2.4 3.8 4.1 7.3 4.9 4.6 4.4 6 6.1 8
4.8 5.9 3.4 2.9 1.7 0.2 0.1 0 0 0.4 0.5 1.3 3.7 13.1 17.3 0.5 12.7 11.8 12.5
19.4 13.5 6.2 3.9 3.5 0.7 0.3 0.1 0.1 0 0 0.1 1.2 3.1 6.1 0.1 12.3 10 7.5
11 11.5 7.3 4.7 2.5 0.2 0.1 0 0.1 0.4 1.1 1.6 3.7 5.9 6.2 2.9 7.8 9.5 10.2
1.5 1.8 2.8 3.4 4 1.5 0.9 0.3 0.3 0.3 0.2 2 9 6.6 8.9 2.3 12 13.8 13.4
1.7 2.5 5.8 7.3 9.3 10.9 6.9 5.1 4.3 4.3 3.9 2.5 0.7 1.4 1.2 1.4 1.1 1 1.2
2.5 2.1 1.3 2.1 1.5 0.7 0.4 0.7 0.8 1.3 1.7 7 8.4 9.5 7.9 5.1 7 6 5.7
1.7 1.8 2.4 2.1 3.1 3.3 3.6 3.2 3.1 2.6 2.2 1.9 1 1.5 1.5 1.4 1.2 1.9 2.3
7.6 7 7 5.8 4 0.6 0.2 0.1 0.1 0 0 0 0.2 0.1 0.2 0.1 0.8 1.7 2.7
0.7 0.8 1.2 1.4 1 1.3 1.5 1.7 2.1 1.8 4.6 4.5 2.1 1.7 1.5 4.6 1.1 0.8 1.2
0.6 0.9 1.4 1.7 2.3 2.9 4 3 2.5 2.8 2.8 1.9 0.7 1.1 1 2 0.3 0.3 0.1
0.3 0.3 0.4 0.2 0.3 0.8 0.9 2 2.6 3.3 3.9 3.3 1.8 1.8 1.7 3.6 1.1 0.8 1
5 1.9 0.9 0.4 0.2 0 0 0 0 0 0.4 0.9 0.4 1.9 3.8 0.2 4.3 4.5 4.5
0 0.1 0.3 0.4 0.9 0.4 0.2 0 0 0 0 0.7 2.7 2.3 3.2 0.8 4.3 5 4.7
0.4 1 1.4 1.7 1.8 2.1 1.7 1.4 1 1.1 0.9 1.3 0.8 1.2 1 0.8 1.7 1.5 1.5
1.3 1.3 1.3 1.4 1.2 0.2 0.1 0.1 0 0.1 0.1 0.3 0.7 2 2.6 0.2 3.1 3.7 4
0.2 0.3 0.3 0.4 0.3 0.9 1.9 1.9 3.8 2.4 2.9 2.1 0.8 0.8 0.6 1.9 0.4 0.4 0.5
0.4 0.8 1.5 1.4 1.7 0.8 0.6 0.5 0.3 0.5 1.3 2.1 2.2 2.1 1.5 3.4 0.6 0.4 0.3
3.9 3.7 2.7 3.7 2.3 0.4 0.2 0.1 0.1 0 0 0 0.1 0.1 0.2 0 1 1.9 1.2
0 0.1 0.5 1 1.7 2.7 2.8 1.9 1.6 1.8 1.4 1.3 0.9 1 0.5 1.2 0 0 0
0 0 0 0.1 0.2 1.6 3.8 2.3 3.2 3 2.4 1.4 0 0.1 0.1 1.2 0 0 0
0.1 0.1 0.3 0.3 0.6 1.3 1.9 1.7 2 2.1 1.5 1.3 0.9 1.1 1 1.7 0.4 0.3 0.1
0 0 0 0 0.1 0.6 1.1 1.2 2.5 2.8 2.8 1.8 0.6 0.5 0.2 3.7 0 0 0
0.1 0.1 0.9 1.1 1.3 0.5 0.4 0.2 0.1 0.3 0.6 1.5 2.6 1.7 1.5 1.3 1.4 1.2 1.1
0.2 0.3 0.5 0.6 0.5 0.1 0 0 0 0 0 0.3 6.4 3.3 2 2.6 0.5 0.2 0.2
0.4 0.4 0.7 0.9 0.8 1.5 1.4 1.4 1.8 1.7 1.2 1.2 0.5 0.5 0.4 1.2 0.4 0.2 0.2
5.7 4.8 1.3 0.9 0.3 0 0 0 0 0 0 0 0 0.4 0.4 0 0.7 1 1.1
2.1 3 3.6 3.6 2.5 0.6 0.1 0.1 0.1 0 0 0 0 0.1 0.1 0 0.2 0.1 0.2
0 0 0 0 0 1.5 3.7 3 4.2 2 0.5 0.3 0.1 0.1 0.1 0.8 0 0 0
1 2 5 7 9 12 14 16 19 21 23 26 28 33 35 37 40 42 44
d
% Read Abundance
20
10
1

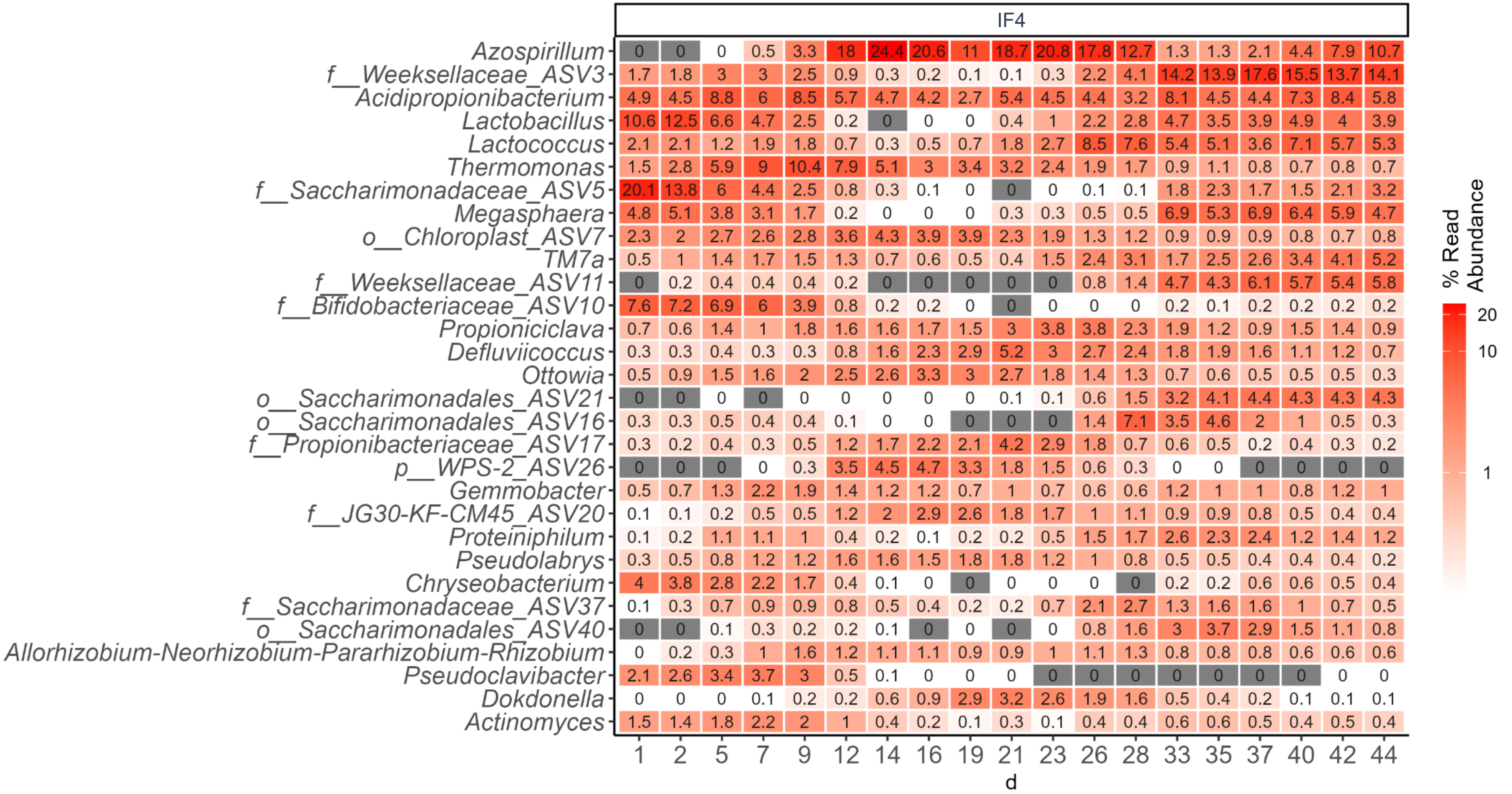


**Figure S4.** Temporal dynamics of the 30 most abundant taxa in replicate reactors IF1–IF4, which were run with inoculum for 44 days. Heatmaps were generated using 16S rRNA gene amplicon (V3–V4) data. The top 30 include genera as well as ASVs that could not be assigned at the genus level but were identified at the family (f_) or order (o_) levels.

**Table S1** – Chemical characteristics of six batches of soybean-processing wastewaters used for microbial community-based SCP production.

| **Soybean wastewater collection batch**[a] | **sCOD** [mg/L] | **sTKN**[b] [mg/L] | **$sPO_4^{3-}$-P** [mg/L] | **$NH_4^+$-N** [mg/L] | **$NO_2^-$-N** [mg/L] | **$NO_3^-$-N** [mg/L] | **TA**[c] [mg/L] | **TSS** [mg/L] | **VSS** [mg/L] | **pH** | **sCOD: sTKN** (C: N)[d] | **sTKN: $sPO_4^{3-}$-P** (N:P)[d] |
|---|---|---|---|---|---|---|---|---|---|---|---|---|
| 1.1 | 3576 | 33.55 | 16.4 | 1.37 | 0.01 | 0.00 | 0 | 220 | 200 | 4.14 | 106.6 | 2.0 |
| 1.2 | 3595 | 36.30 | 16.7 | 1.24 | 0.01 | 0.00 | 0 | 220 | 180 | 4.14 | 99.0 | 2.2 |
| 2.1 | 4087 | 42 | 20.0 | 1.36 | 0.01 | 0.00 | 25 | 360 | 340 | 4.16 | 97.5 | 2.1 |
| 2.2 | 2820 | 26 | 12.1 | 0.91 | 0.00 | 0.00 | 4 | 200 | 100 | 4.16 | 108.1 | 2.2 |
| 3.1 | 2839 | 24 | 14.9 | 0.93 | 0.00 | 0.03 | 0 | 140 | 140 | 4.11 | 117.6 | 1.6 |
| 3.2 | 2575 | 21 | 13.4 | 0.88 | 0.00 | 0.03 | 0 | 120 | 120 | 4.11 | 120.4 | 1.6 |
| 3.3 | 4106 | 31 | 23.7 | 1.77 | 0.00 | 0.01 | 16 | 100 | 100 | 4.11 | 133.4 | 1.3 |
| 3.4 | 8751 | 95 | 62.8 | 2.63 | 0.00 | 0.05 | 131 | 260 | 200 | 4.11 | 91.8 | 1.5 |
| 4.1 | 8245 | 62 | 48.1 | 5.13 | 0.02 | 0.08 | 0 | 400 | 400 | 3.81 | 134.0 | 1.3 |
| 4.2 | 8763 | 76 | 58.6 | 4.64 | 0.03 | 0.02 | 15 | 500 | 500 | 3.81 | 116.0 | 1.3 |
| 5.1 | 8362 | 61 | 49.6 | 6.50 | 0.00 | 0.04 | 0 | 300 | 300 | 3.82 | 137.5 | 1.2 |
| 5.2 | 7674 | 62 | 48.3 | 6.57 | 0.00 | 0.04 | 0 | 360 | 360 | 3.82 | 123.2 | 1.3 |
| 6.1 | 10132 | 88 | 67.9 | 10.77 | 0.02 | 0.06 | 0 | 400 | 380 | 3.94 | 115.7 | 1.3 |
| 6.2 | 11370 | 109 | 76.9 | 10.98 | 0.00 | 0.05 | 0 | 360 | 360 | 3.94 | 104.0 | 1.4 |
| Average[e] (s.d.m.) | 6207 (2984) | 55 (28) | 38 (22) | 3.98 (3.44) | 0.01 (0.01) | 0.03 (0.02) | 14 (33) | 281 (116) | 263 (125) | 4.01 (0.14) | 115 (14) | 1.6 (0.4) |

[a] A total of fourteen 20-L carboys collected at six different time points over two months. The soybean wastewater collection batches were numbered from 1 to 6, with Batch 3 consisting of four carboys (3.1, 3.2, 3.3, 3.4), while all others consisted two carboys.

[b] sTKN was calculated using the value of sTN, nitrate, and nitrite: sTKN = sTN – nitrate - nitrite.

[c] Total alkalinity expressed as calcium carbonate

[d] Value calculated from the measured chemical data.

[e] Average values of influent wastewater batches used, including standard deviation of the mean (s.d.m.) in parentheses.

**Table S2.** Chemical characteristics of SCP-producing bioreactor effluent reused in downstream heat treatment of microbial protein.

| **Heat treatment phase** | **sCOD** [mg/L] | **sTOC** [mg/L] | **sTN** [mg/L] | **sTKN** [mg/L] | **TA** [mg/L] | **$sPO_4^{3-}$-P** [mg/L] | **$NH_4^+$-N** [mg/L] | **$NO_2^-$-N** [mg/L] | **$NO_3^-$-N** [mg/L] | **$Cl^-$** [mg/L] | **$SO_4^{2-}$-S** [mg/L] | **pH** |
|---|---|---|---|---|---|---|---|---|---|---|---|---|
| 1 | 300 | 28 | 29.2 | 22.4 | 316 | 0.7 | 0 | 0 | 6.8 | 2.0 | 2.4 | 7.3 |
| 2 | 84 | 28 | 1.7 | 1.2 | 494 | 13.1 | 0 | 0 | 0.5 | 23.2 | 5.5 | 7.5 |

sCOD: Soluble chemical oxygen demand
sTOC: Soluble total organic carbon
sTN: Soluble total nitrogen
sTKN: Soluble total kjeldahl nitrogen
TA: Total alkalinity expressed as calcium carbonate
$sPO_4^{3-}$-P: Soluble phosphate expressed as phosphorus
$NH_4^+$-N: Ammonium expressed as nitrogen
$NO_2^-$-N: Nitrite expressed as nitrogen
$NO_3^-$-N: Nitrate expressed as nitrogen
$Cl^-$: Chloride
$SO4^{2-}$-S: Sulphate expressed as sulphur

**Table S3 -** Chemical characteristics of bioreactor effluent and influent.

| Reactor | Day | sCOD_effluent[b] [mg/L] | sCOD_feed[c] [mg/L] | sTN_effluent[b] [mg/L] | sTN_feed[c] [mg/L] |
|---|---|---|---|---|---|
| IF1[a] | 2 | 438 | 3576 | | |
| | 5 | 413 | 3576 | | |
| | 7 | 209 | 4087 | 4 | 42 |
| | 9 | 168 | 4087 | 2 | 42 |
| | 12 | 153 | 4087 | 10 | 42 |
| | 14 | 231 | 2839 | 9 | 24 |
| | 16 | 264 | 2575 | 9 | 21 |
| | 19 | 314 | 2575 | 10 | 21 |
| | 21 | 1170 | 8751 | 12 | 95 |
| | 23 | 1357 | 8751 | 16 | 95 |
| | 26 | 1480 | 8763 | 15 | 76 |
| | 28 | 2702 | 8245 | 16 | 62 |
| | 33 | 2657 | 7674 | 19 | 62 |
| | 35 | 2111 | 8362 | 23 | 61 |
| | 37 | 556 | 8362 | 7 | 61 |
| | 40 | 385 | 11370 | 5 | 109 |
| | 42 | 404 | 10132 | | |
| | 44 | 56 | 10132 | | |
| IF2[a] | 2 | 520 | 3576 | | |
| | 5 | 500 | 3576 | | |
| | 7 | 231 | 4087 | 3 | 42 |
| | 9 | 147 | 4087 | 2 | 42 |
| | 12 | 157 | 4087 | 9 | 42 |
| | 14 | 249 | 2839 | 8 | 24 |
| | 16 | 251 | 2575 | 11 | 21 |
| | 19 | 243 | 2575 | 8 | 21 |
| | 21 | 1046 | 8751 | 9 | 95 |
| | 23 | 1558 | 8751 | 16 | 95 |
| | 26 | 952 | 8763 | 13 | 76 |
| | 28 | 1692 | 8245 | 11 | 62 |
| | 33 | 3191 | 7674 | 19 | 62 |
| | 35 | 1323 | 8362 | 15 | 61 |
| | 37 | 378 | 8362 | 6 | 61 |
| | 40 | 709 | 11370 | 6 | 109 |
| | 42 | 1396 | 10132 | | |
| | 44 | 628 | 10132 | | |
| IF3[a] | 2 | 411 | 3576 | | |
| | 5 | 739 | 3595 | | |
| | 7 | 182 | 2820 | 3 | 26 |
| | 9 | 153 | 2820 | 2 | 26 |
| | 12 | 186 | 2820 | 10 | 26 |
| | 14 | 223 | 2575 | 8 | 21 |
| | 16 | 230 | 2839 | 5 | 24 |
| | 19 | 196 | 2839 | 5 | 24 |
| | 21 | 482 | 4106 | 7 | 31 |
| | 23 | 1937 | 8751 | 16 | 95 |
| | 26 | 1656 | 8245 | 8 | 62 |
| | 28 | 2839 | 8763 | 16 | 76 |
| | 33 | 3391 | 8362 | | |
| | 35 | 3773 | 7674 | 21 | 62 |

**Table S3.** *(cont.).*

| Reactor | Day | sCOD_effluent[a] [mg/L] | sCOD_feed[b] [mg/L] | sTN_effluent[a] [mg/L] | sTN_feed[b] [mg/L] |
|---|---|---|---|---|---|
| IF3[a] | 37 | 3357 | 7674 | 21 | 62 |
| | 40 | 4946 | 10132 | 27 | 88 |
| | 42 | 7149 | 11370 | | |
| | 44 | 7123 | 11370 | | |
| IF4[a] | 2 | 446 | 3576 | | |
| | 5 | 701 | 3595 | | |
| | 7 | 176 | 2820 | 3 | 26 |
| | 9 | 156 | 2820 | 3 | 26 |
| | 12 | 186 | 2820 | 8 | 26 |
| | 14 | 208 | 2575 | 8 | 21 |
| | 16 | 225 | 2839 | 8 | 24 |
| | 19 | 257 | 2839 | 10 | 24 |
| | 21 | 336 | 4106 | 6 | 31 |
| | 23 | 1095 | 8751 | 14 | 95 |
| | 26 | 1851 | 8245 | 12 | 62 |
| | 28 | 2196 | 8763 | 14 | 76 |
| | 33 | 2245 | 8362 | 16 | 61 |
| | 35 | 1451 | 7674 | 16 | 62 |
| | 37 | 771 | 7674 | 10 | 62 |
| | 40 | 1302 | 10132 | 13 | 88 |
| | 42 | 1233 | 11370 | | |
| | 44 | 360 | 11370 | | |

[a] Four replicate reactors (IF1-4) were run for 44 d with a microbial inoculum under identical conditions.

[b] Effluent discharged after 1 h of settling, before feeding phase.

[c] Influent wastewater used right after effluent discharge.
